# Microbiology Galaxy Lab: The first community-driven gateway for reproducible and FAIR analysis of microbial data

**DOI:** 10.1101/2024.12.23.629682

**Authors:** Engy Nasr, Nikos Pechlivanis, Nikolaos Strepis, Pierre Amato, Matthias Bernt, Anshu Bhardwaj, Daniel Blankenberg, Daniela Brites, Fabio Cumbo, Katherine T Do, Emanuele Ferrari, Timothy J. Griffin, Björn Grüning, Saskia Hiltemann, Cameron J Hyde, Pratik Jagtap, Subina Mehta, Kimberly L. Métris, Saim Momin, Tiffanie M. Nelson, Asime Oba, Christina Pavloudi, Raphaëlle Péguilhan, Gareth R Price, Fotis Psomopoulos, Nedeljka Rosic, Michael C. Schatz, Valerie Claudia Schiml, Cléa Siguret, Nicola Soranzo, Andrew Stubbs, Peter van Heusden, Mustafa Vohra, microGalaxy Community, Paul Zierep, Bérénice Batut

## Abstract

The explosion of microbial omics data has outpaced the ability of many researchers to analyze it, with complex tools and limited computational resources creating barriers to discovery. To address this gap, we present the Microbiology Galaxy Lab: a free, globally accessible, community-supported platform that combines state-of-the-art analytical power with user-friendly accessibility. Supported by the Galaxy and global microbiology communities, this platform integrates over 315 tool suites and 115 curated workflows, enabling comprehensive metabarcoding, (meta)genomic, (meta)transcriptomic, and (meta)proteomic data analysis within a FAIR-aligned environment. It also supports research in the health and infectious disease sectors, as well as in environmental microbiology. The platform’s utility is exemplified through various use cases, including antimicrobial resistance tracking, biomarker prediction, microbiome classification, and functional annotation of key microbes. Built on reproducibility and community engagement, it supports creation, sharing, and updating of best-practice workflows. Over 35 tutorials and learning paths empower scientists, fostering an ecosystem that keeps resources at the forefront of microbial science. The Microbiology Galaxy Lab enables collective analysis, democratising research, thereby accelerating discovery across the global microbiology community (microbiology.usegalaxy.org, microbiology.usegalaxy.eu, microbiology.usegalaxy.org.au, microbiology.usegalaxy.fr).

## Introduction

Over the past two decades, microbiology has experienced a transformative shift due to advances in molecular biology and high-throughput sequencing technologies^1,2^. These innovations have enhanced our ability to analyse microbes in human and animal health, as well as in ecosystem contexts. To derive meaningful insights into microbial pathogens, community compositions, and functional traits, it is essential to employ scalable bioinformatics tools that can process and interpret massive and diverse datasets. This constant need for advanced analytical approaches, such as gut microbiome analysis, efficient detection of resistant microbes or pathogens, and microbial genome assembly and annotation, highlights a critical bottleneck: many microbiologists lack the skills for these intricate analyses. This challenge is further magnified by microbiology bioinformatics demands, requiring appropriate bioinformatic tools, computational resources, and access to appropriate reference databases^3,4^. Closing the bioinformatics knowledge gap and making powerful analytical pipelines widely available is therefore essential, and the microbiology field is increasingly focusing on open, standardised computing environments^5^. Effective solutions must be robust, accurate, reproducible, and adhere to the FAIR (Findable, Accessible, Interoperable, Reusable) principles^6^ to reduce complexity in data organisation, annotation, and tool compatibility, thereby supporting wider advanced methods adoption.

A large amount of microbial data and metadata is produced by healthcare and research centers, providing a crucial foundation for understanding pathogen spread and working towards pandemic preparedness^7^. Public health systems have benefited from incorporating whole genome sequencing (WGS) in routine diagnostic or surveillance studies^7^. However, WGS data is often not fully utilised due to limitations in bioinformatics expertise and infrastructure^8^. In particular, global disparities in methodologies and access to computational infrastructure reduce researchers’ ability to perform large-scale microbial analyses^3^.

Assessing data quality, assembling genomes, metagenomes, and plasmids, and detecting resistance and virulence traits are essential steps for understanding complex infections and pathogen spread. When combined with other omics—such as proteomics—these analyses can reveal functional pathogen mechanisms for future clinical use. Because pathogen–host interactions are inherently complex, a systems-biology approach that integrates multiple omics layers is needed^9,10^ to capture taxonomic, functional, and expression-level information and to untangle infection mechanisms.

Unlocking insights into microbial biodiversity and ecosystem function requires reproducible, scalable workflows. Environmental microbiology research increasingly uses multi-omic technologies, integrating data from (meta)genomics, (meta)transcriptomics, (meta)proteomics, and metabolomics to explore microbial community composition, functional traits, and responses to stressors like pollution and climate change^11–13^. These approaches are essential for understanding microbial roles in nutrient cycling, carbon dynamics, and environmental resilience^12^. Data integration from diverse environments, such as wastewater, plant, and atmosphere sources, further advances our understanding of disease and infection emergence, as demonstrated by multi-omic data analyses to track resistant species^14,15^.

Interdisciplinary frameworks like One Health^16^ benefit from data integration and analysis supporting microbiology research across public health, agriculture, and environmental systems.

To address the need for flexible and accessible microbial bioinformatics, several platforms have been developed (**Extended Data Table 1**). While these platforms excel in specific domains, many are limited to certain ‘omics modalities, lack interoperability, or function as “black boxes”, reducing user transparency and control^17^. Additionally, limited computational expertise remains a barrier to widespread adoption, underscoring the need for open, integrated, and user-friendly solutions.

The growing demand for a user-friendly platform that can handle data-intensive biomedical analysis and computing needs led to the development of Galaxy^18^. Galaxy is an open, web-based, and widely adopted platform for FAIR-compliant, reproducible research suitable for robust bioinformatics analyses in many domains, including microbiology. A survey of Galaxy users working in microbial research revealed that they primarily use the platform for bacterial genomics and metagenomics, with a growing interest in functional analysis, gene identification, and microbial genome assembly (**Supplementary Table 1**). The survey results also highlighted the emerging need in areas such as metaproteomics and resistance profiling, and challenges in limited user experience and technical hurdles, underscoring the need for accessible tools and community-driven support. An evolution of these identified challenges led to the formation of the microGalaxy community, which consists of practitioners from microbiological research, including microbiologists and bioinformaticians. The survey results and the ongoing exchange within the microGalaxy community led to the development of the Microbiology Galaxy Lab, a collective open-source Galaxy-based platform for streamlined and reproducible microbial data analysis freely available on several Galaxy servers (microbiology.usegalaxy.org, microbiology.usegalaxy.eu, microbiology.usegalaxy.org.au, microbiology.usegalaxy.fr).

The Microbiology Galaxy Lab builds on Galaxy’s foundation to address the microbial bioinformatics challenges and create analysis opportunities by offering: (i) access to a wide range of critically reviewed software tools and workflows, (ii) a user-friendly interface, (iii) a powerful workflow management system including a graphical editor, (iv) free access to public computational resources, (v) integration with training resources to support users with varying bioinformatics expertise, and (vi) automation, integration, and advanced data management through command line tools and comprehensive APIs. By leveraging these features, the Microbiology Galaxy Lab provides a comprehensive framework supporting the global microbiology community in overcoming data and computational barriers. The use of Microbiology Galaxy Lab as a versatile and community-driven infrastructure significantly enhances health and disease, and ecosystems and biodiversity research. By integrating microbial and phenotypic data analyses, such as antibiotic resistance profiles or ecological traits, it fosters interdisciplinary research and promotes standardised, reproducible approaches. This integration advances microbial data analysis and deepens our understanding of microbial and ecosystem diversity, including their roles in health and global environmental change.

## Results

### Galaxy adoption for microbial research: community trends and insights

To indicate the value of user-friendly implementations for complex computational workflows, a dual-pronged analysis was conducted: a text-mining review of published literature leveraging the Galaxy platform for microbial analysis, complemented by a targeted microbial community survey. Our findings reveal a consistent and upward trend in Galaxy utilisation since 2005, with a pronounced increase in microbial applications. Notably, between 2020 and 2025, 4,891 publications relied on the Galaxy platform for their analysis, of which 1,911 were specifically focused on microbial computational processes **(Fig. 1A**). Among these, 559 studies (29.3%) addressed health and disease applications, 364 (19.0%) focused on ecosystems and biodiversity, and a significant portion, 988 publications (51.7%), bridged both domains, reflecting the growing interdisciplinarity of microbial research **(Extended Data Fig. 1**). Within this subset, (meta)genomics, a computationally intensive field, was the most prevalent application, followed by metabarcoding and metatranscriptomics. This focus on meta-omics studies reflects the inherent complexity of microbiological data and diversity. Key analytical functions identified in the literature included taxonomic classification, functional analysis, and antimicrobial resistance (AMR) gene profiling, with target taxonomic groups spanning bacteria, pathogens, microbiomes, and viruses (**Extended Data Fig. 1**).

**Figure 1:**
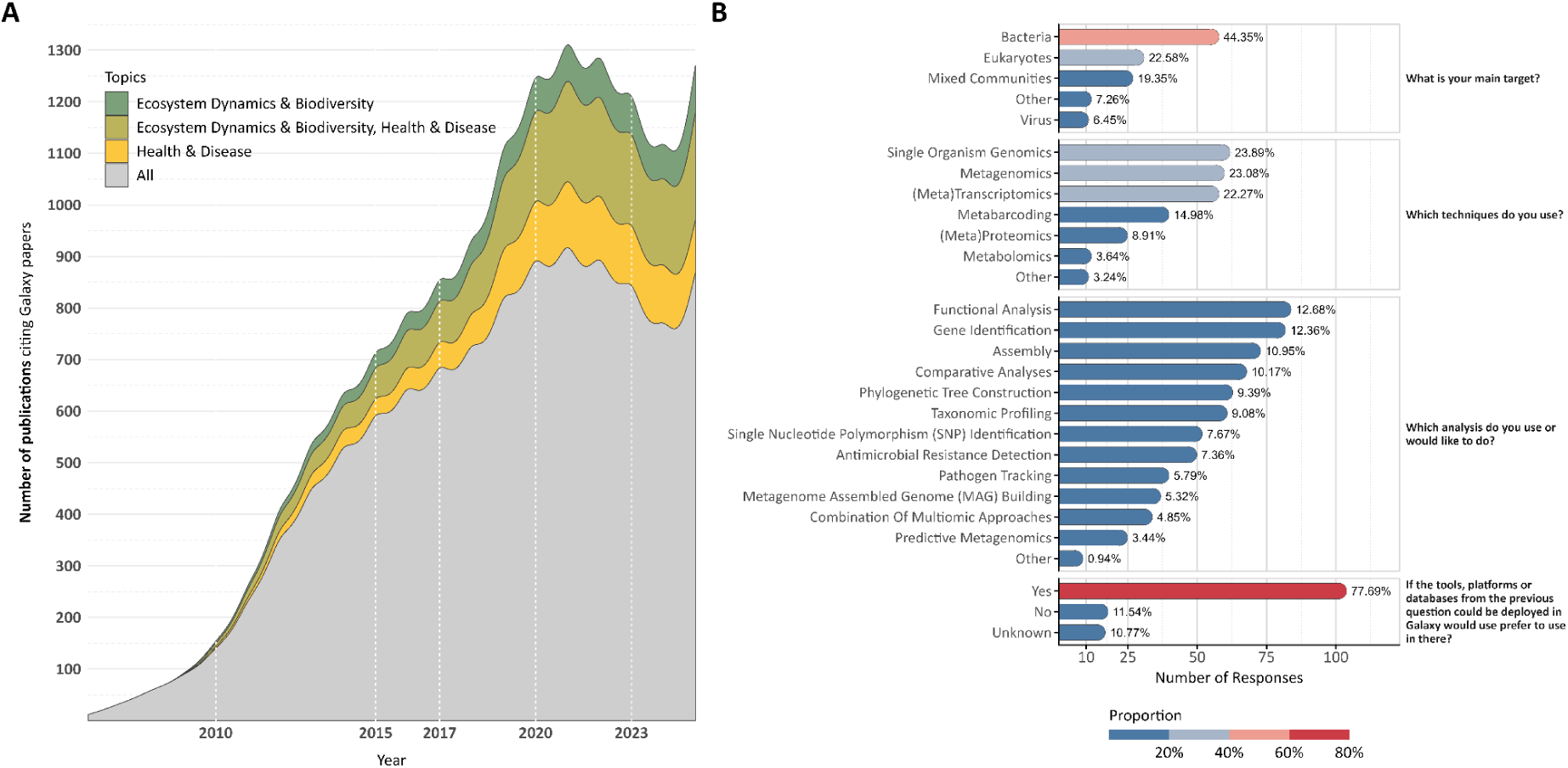
Citation trends in Galaxy publications and microbial research community survey. The citations were extracted from the Galaxy Project’s Google Scholar profile, and additional details were retrieved using Semantic Search. (A) Annual publication trends show the total number of citations (gray) alongside those specific to microbial research divided into health and disease (yellow), ecosystem dynamics and biodiversity (green), and for both (olive yellow). (B) Survey results from the microbiology research community (March–September 2023). Anonymous responses highlighting the main research targets, techniques, and analyses used or desired by microbial researchers. The figure also reports challenges faced by users who rarely or never use Galaxy, and preferences regarding tool deployment in Galaxy. Percentages were computed independently for each question based on the total number of responses available for that question. Additionally, some questions, such as “*Which analyses do you use or would you like to do?*” allowed multiple responses, meaning percentages may sum to more than 100%.

Furthermore, to gather community feedback, a survey was distributed to Galaxy users and shared with the microbial community beyond Galaxy. Questions were designed to assess current microbial workflow applications and identify areas for enhancement (**Supplementary Document 1**). The survey provides a granular look into the priorities and practices of the community of microbial bioinformaticians and researchers. The results show a clear focus on microbial multidisciplinary methodologies, with bacteria being the main target organism for a substantial proportion of respondents (44.4%, **Fig. 1B, Supplementary Table 1**). The top three techniques were split between single-organism genomics (23.9%), metagenomics (23.1%), and (meta)transcriptomics (22.3%), indicating diverse but clustered popular approaches. Regarding specific computational tasks, functional analysis (12.7%) is the most frequently used or desired analysis, followed closely by gene identification (12.4%) and genome assembly (11.0%). A Galaxy key strength is its modularity, allowing researchers to mix and integrate various tools and workflows for their specific research aims. As confirmed by user surveys, accompanying tools and workflows with tutorials significantly increases their adoption.

The most insightful result regarding microbiology computational analysis is that the primary barrier to its adoption is lack of experience (51.9%) and technical difficulties (25.9%), and not lack of utility (**Supplementary Table 1**). This is underscored by the fact that an overwhelming number of respondents (77.7%) would prefer to use Galaxy if it integrated other tools and databases they needed (**Fig. 1B**). This presents a clear opportunity for a focused platform leveraging microbial tools to enhance the user experience. The survey reveals that the platform’s user base is currently concentrated in Europe (31.7%) and Asia (27.9%, **Supplementary Table 1)**; thus, its most transformative potential lies in serving researchers in developing countries. A microbiology-focused Galaxy platform would help these scientists to analyse their valuable datasets with computationally complex methods, bypassing their local infrastructure limitations and barriers to promote scientific self-reliance.

### Community-curated and ready-to-use tools and workflows on Microbiology Galaxy Lab

Over the past five years, tools on the Microbiology Galaxy Lab have supported over 25 million analyses, either as standalone applications or within workflows, underscoring their broad utility in the microbial research community (**Fig. 2A, Supplementary Table 2**). Metrics on tool usage provide insight into user engagement and platform impact: a strong positive correlation is observed between the number of unique users and individual tool execution frequency (**Fig. 2B, C**), suggesting widespread adoption of key computational functions.

**Figure 2:**
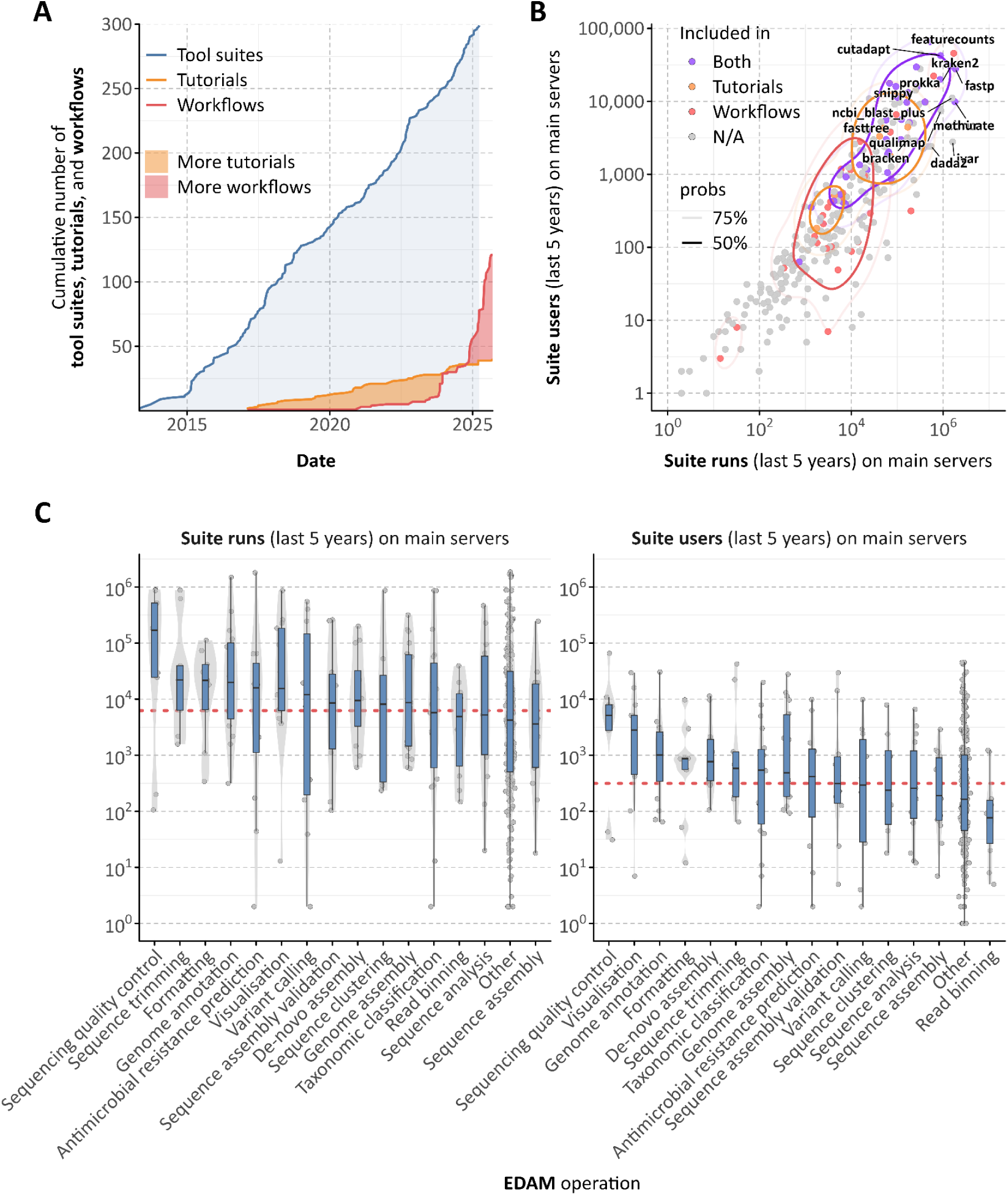
Growth and usability of microbiology-related tool suites, training materials/tutorials, and workflows within the Galaxy ecosystem. (A) Cumulative number of microbiology-related tool suites, tutorials, and workflows added to Galaxy over time, based on the date of the first commit. (B) Scatter plot showing the usage (number of runs and number of users) of microbiology-related tool suites, tutorials, and workflows over the past five years across all Galaxy main servers. (C) Box plot showing the usage of microbiology-related tool suites over the past five years across all Galaxy main servers, categorised by their EDAM operations.

The Microbiology Galaxy Lab offers powerful and diverse tool suites designed for large-scale microbial data analysis. It includes 884 community-curated tools derived from 318 tool suites within the broader Galaxy ecosystem, specifically tailored for a wide range of microbial applications (**Fig. 3, Supplementary Table 2, Extended Data Fig. 2**). Through these tools, users have access to over 170 integrated reference genomes and 11 terabytes of reference data, delivered through CernVM-FS^19^, a distributed network file system optimised for scientific software and data. These reference datasets support essential microbial bioinformatics tasks such as taxonomic classification and functional annotation.

**Figure 3:**
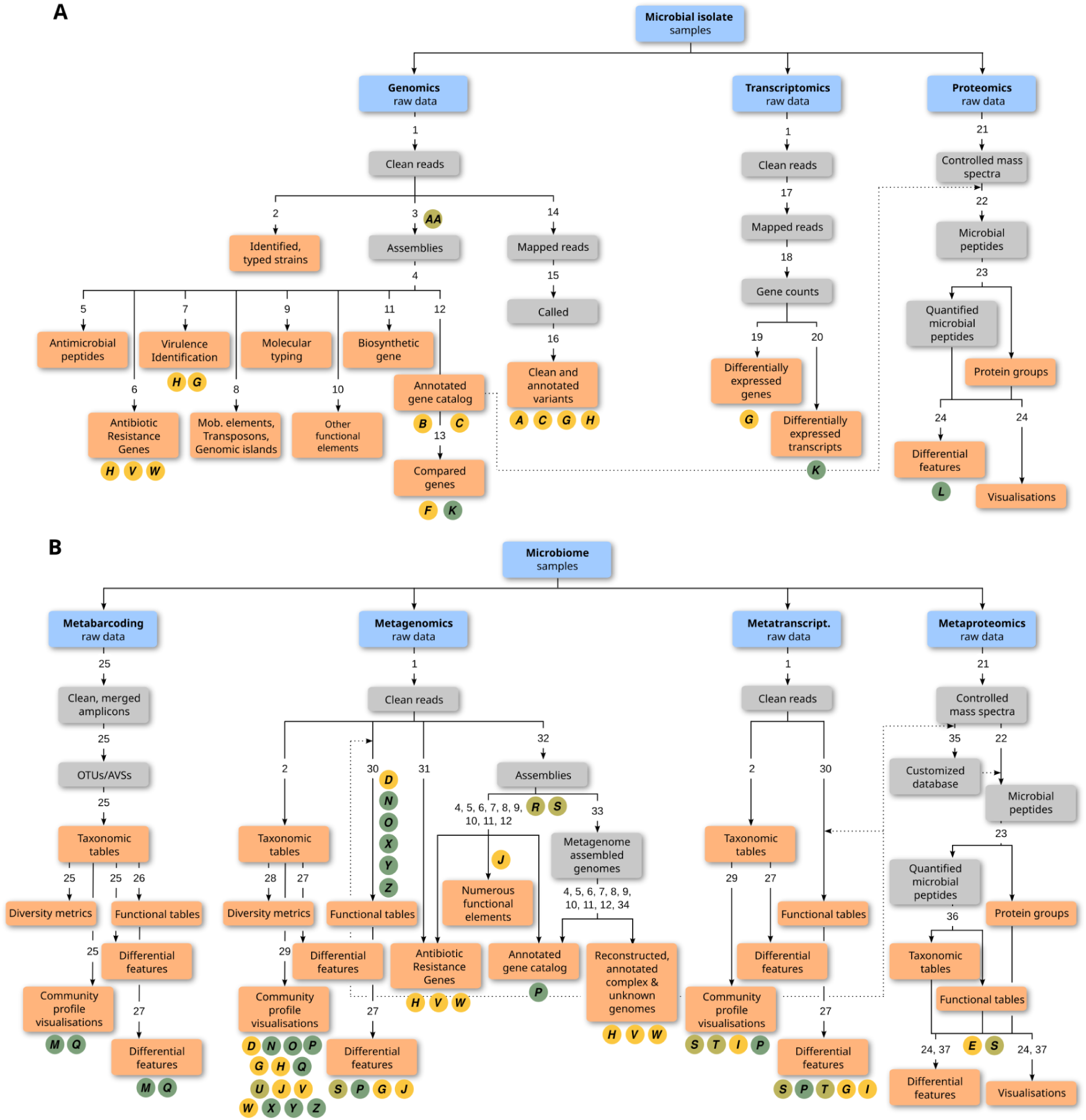
Overview of microbial data analysis tasks with corresponding tool suites and potential analyses available on the Microbiology Galaxy Lab. Analysis tasks, categorised by data type, for (A) microbial isolate samples, including genomics, transcriptomics, and proteomics, and (B) microbiome samples, encompassing metabarcoding, metagenomics, metatranscriptomics, and metaproteomics. Each task progresses from raw data (blue) to intermediate (gray) and final outputs (coral). Circled letters indicate use cases, as listed in **Supplementary Table 3**. They are color-coded by domain: health and disease (yellow), ecosystem dynamics and biodiversity (green), and for both (olive yellow). Numbers on the branches correspond to tool suites that can be used for generating the outputs below, and they are detailed in **Supplementary Table 4.**

The platform also supports interactive tools enabling personalised microbial data exploration. Examples include Pavian^20^, Phinch^21^, Shiny Phyloseq^22^, and Apollo^23^, facilitating results’ real-time inspection and curation within the platform. Additionally, general-purpose tools such as Jupyter Notebooks (<u>RRID:SCR_018315</u>) and RStudio (<u>RRID:SCR_000432</u>) enhance interactivity and reproducibility, supporting modular and shareable analyses.

Complementing its toolset, the Microbiology Galaxy Lab hosts 117 curated, ready-to-use workflows **(Supplementary Table 5, Extended Data Fig. 3**), streamlining complex analyses into reproducible, user-friendly processes. These workflows reflect a broad consensus within the microGalaxy community and are developed by domain experts in microbiology and bioinformatics. Forty-two workflows are published in public repositories such as WorkflowHub^24^ and Dockstore^25^ and can be directly executed within the Microbiology Galaxy Lab or installed on any Galaxy instance. The remaining workflows are shared across Galaxy servers under the *microgalaxy* tag, enabling direct discovery and reuse. To make full use of these workflows and facilitate customisation, a key strength of the platform is its graphical interface, allowing users to create, customise, and manage workflows interactively, significantly lowering the technical barrier to designing and executing complex analyses.

Although not directly transferable, selected workflows can be conceptually adapted or re-implemented in other systems such as Nextflow^26^ and Snakemake^27^, providing flexibility for users who wish to deploy them within alternative workflow engines. To support users with diverse bioinformatics expertise, the platform includes training materials and tutorials for using and interpreting microbial workflows, detailed in the following subsection. The tools, workflows, and associated training materials are continuously maintained and expanded to meet emerging needs in microbial research (**Fig. 2A**).

### Tools and workflows supporting applications in health, disease, and ecosystem research

Tools and workflows in the Microbiology Galaxy Lab offer lots of diverse methodologies **(Fig. 3**) to support a wide range of applications in health, disease, and ecosystem research (**Fig. 4, Supplementary Table 3)**.

**Figure 4:**
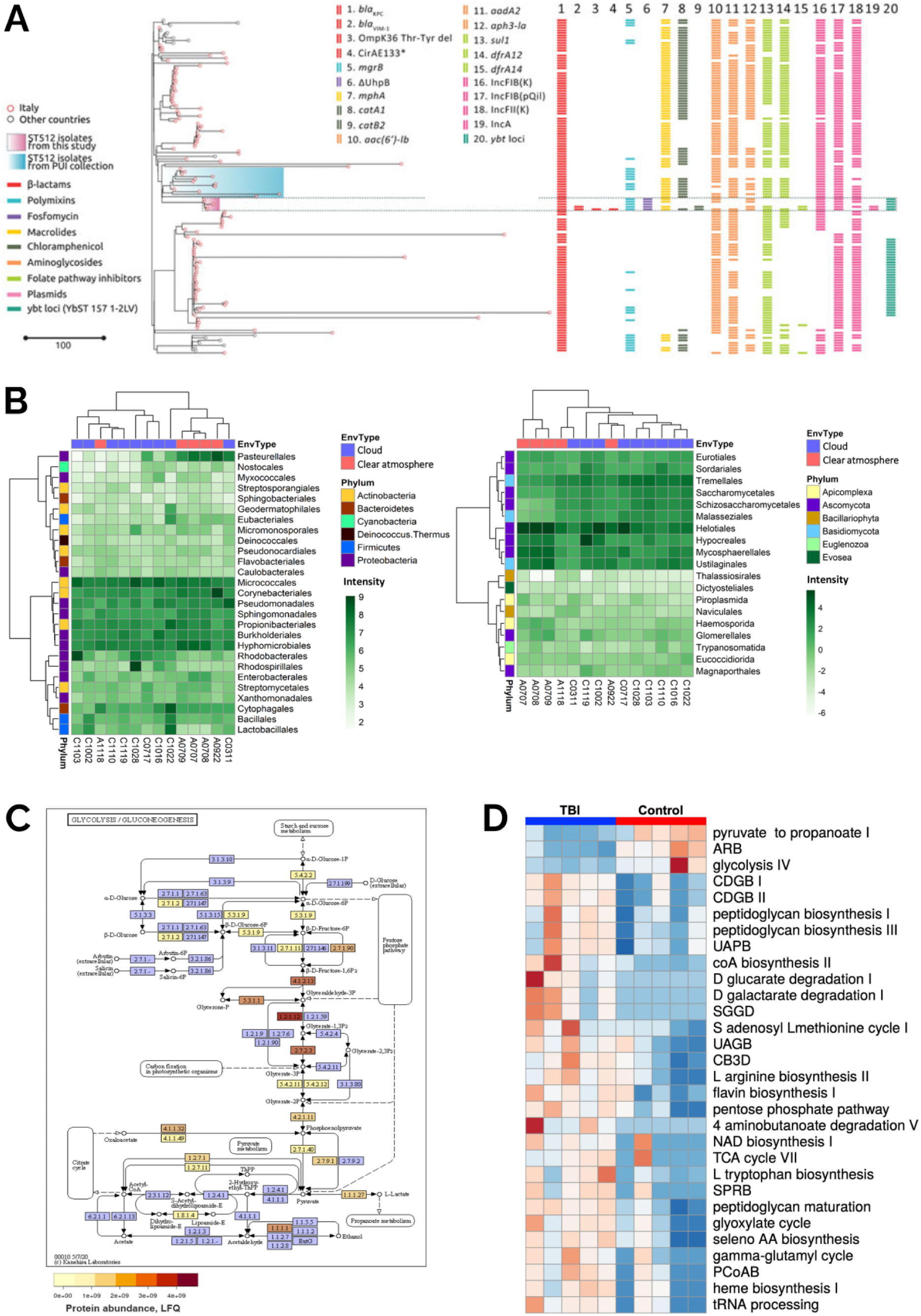
Example of applications as tools and workflows available in the Microbiology Galaxy Lab in health, disease, and ecosystem research. (A) Phylogenetic and genetic analysis of *Klebsiella pneumoniae* ST512, illustrating its genotypic evolution under treatment with ceftazidime/avibactam, meropenem/vaborbactam, and cefiderocol (Figure 2 in Arcari et al 2023^28^). (B) Comparative distribution of dominant bacterial and eukaryotic orders in cloudy versus clear-atmosphere metagenomes, with hierarchical clustering (Figure 1 in Péguilhan et al. 2025^29^). (C) Glycolysis/Gluconeogenesis pathway of a *Tissierellia*-class bacterium, built as a Metagenomics Assembled Genome from metagenomics data and annotated with metaproteomic abundance (Figure 3 in Schiml et al 2023^30^). (D) Heatmap of 30 differentially expressed biochemical pathways in traumatic brain injury (TBI) versus control fecal metatranscriptomes (Figure 3 in Pyles et al 2024^31^).

#### Health and disease

In the health and disease context, Microbiology Galaxy Lab microbial bioinformatics workflows are indispensable for elucidating microbial pathogenesis and host-microbe interaction mechanisms. A primary application is using toolsets to analyse how pathogens alter their gene composition, expression, and protein production to survive within the host environment, resist the immune system, or develop antibiotic resistance^32^. Workflows analogous to metagenomic assembly are critical for studying biofilm formation and cell sensing, which are key virulence strategies allowing bacteria to establish chronic infections and resist treatment^30^. All these computational microbiology analyses are built upon homogeneity checks to ensure data from different patient samples are comparable, and technology suites to validate diagnostic and sequencing methods.

AMR genes pose a significant threat to healthcare^33^, making the comprehensive analysis of their presence in both clinical and environmental samples essential for controlling their spread and treating severe infections. With the growing volume of clinical sequencing for routine diagnostics, there is a critical need for intuitive analysis platforms accessible to non-specialists that support AMR analysis, including genotype identification and its correlation with carrying plasmids through plasmid identification. ABRomics (https://www.abromics.fr/), a platform for AMR research and surveillance, is built on Microbiology Galaxy Lab with workflows developed in consultation with the microGalaxy community of experts. Workflows encompass a range of stages from sample sequencing quality assessment, read cleaning and trimming, to taxonomic assignment, genome assembly, genome annotation, and AMR gene identification. As AMR genes are spread through mobile genetic elements, workflows that can screen genomes and identify plasmids, integrons, and insertion sequence elements were developed to expand ABRomics and provide more complete information regarding AMR analysis. Tools and workflows for defining AMR genotype are multiple, offering choices and often generating diverse results due to different reference databases, algorithms, or parameters. BenchAMRking^34^ was developed to demonstrate challenges in AMR identification and issues with AMR gene ontologies, underscoring the necessity for greater standardisation in AMR research. Furthermore, BenchAMRking incorporates an ISO-certified AMR identification^35^ tool that generates a clinical report, making it suitable for implementation in healthcare settings. To further bridge the gap between genotype and phenotype, Galaxy-ASIST^36^ integrates antimicrobial susceptibility testing (AST) data with genomic AMR profiles, supporting harmonised interpretation and clinical reporting aligned with global standards^36^. The versatility of Microbiology Galaxy Lab enables the study of diverse microbial taxa, as illustrated by Merdan et al. 2022^37^ who identified drug-resistance mutations in *Candida glabrata* conferring resistance to azole and amphotericin B. A related challenge is complex microbiome data analysis, demanding substantial computational power and specialised expertise to extract AMR genes and link them to their microbial hosts. To address this, dedicated workflows are being developed, especially by the ABRomics team.

In parallel, clinical metaproteomics is emerging as a powerful approach in microbiome research, offering insights into host-microbe interactions underlying human disease. However, challenges persist, primarily in characterising microbial proteins present in low abundance relative to host proteins. Researchers also face difficulties in handling large protein sequence databases for peptide and protein identification from mass spectrometry data, as well as in performing taxonomic and functional assignments and statistical analysis to identify differentially abundant peptides. Galaxy workflows tackle these challenges, supporting studies on COVID-19, cystic fibrosis, and ovarian cancer biomarker discovery^32,38,39^.

Complementing these efforts, PathoGFAIR^40^ provides a standardised, FAIR-compliant framework within Galaxy for foodborne pathogen detection and characterisation from high-throughput sequencing data. PathoGFAIR integrates end-to-end, curated workflows for quality control, taxonomic profiling, antimicrobial resistance, and virulence factor identification. By enforcing interoperability and rich metadata annotation, PathoGFAIR facilitates reproducible analyses and cross-institutional harmonisation in pathogen surveillance and outbreak investigations. Its public availability via Galaxy servers ensures broad accessibility, supporting both research and routine public health applications within a One Health paradigm.

Applying machine learning and artificial intelligence to microbiome research is a growing interest area, successfully integrated into Galaxy, allowing researchers to perform more efficient analyses. Cumbo et al. 2024^41^ used AI-driven tools to identify potential microbial biomarkers for colorectal cancer by analysing public metagenomic samples from patients in a case/control scenario.

#### Ecosystem dynamics and biodiversity

Understanding microbial ecosystem dynamics is fundamental to the planet’s survival as the former drive critical processes like nutrient cycling, carbon sequestration, and decomposition^42,43^. Microbial taxonomic classification is primarily achieved through metabarcoding, metagenomics, and whole-genome sequencing of isolated strains. However, this presents major challenges, e.g., for metagenomics: shotgun sequencing data volume, assembling genomes from mixed reads (binning), computational complexity, and the need for deep, accurate data to resolve strain-level differences^44,45^. Overcoming these hurdles requires integrated platforms with high computational capacity. By making processes reproducible and accessible, such platforms empower the research community to conduct large-scale data analysis to answer their research objectives efficiently and easily.

The Microbiology Galaxy Lab contains workflows for short and/or long-read sequencing to classify ecosystem members and interrogate and decipher community dynamics. Long-read sequencing in combination with short reads was employed to identify *Trichoderma* species strains, demonstrating suitability for hybrid assemblies and correct classification even at the strain level^46^. As mentioned before, microbiome analyses are often associated with 16S rRNA amplicon sequencing; amplicon data were used to explore root microbiomes, and species classification tools available in the Lab allowed for the identification of bacterial species essential for promoting plant growth^47^. Workflows available in Microbiology Galaxy Lab also enhance microbial sequencing analysis by advancing existing pipelines for accurate taxonomic species classification or further improved downstream analysis. In collaboration with EMBL-EBI, the MGnify^5^ amplicon analysis pipeline has been adapted into a customisable Galaxy workflow^48^, benefiting the standardisation of practices across platforms.

Microbiology Galaxy Lab supports participatory microbiome classification and education. The BeerDEcoded project by the Street Science Community (https://streetscience.community/), engaged diverse participants to analyse microbiomes from beer samples. The platform’s user-friendly interface and accessible workflows have enabled high school students to explore the classification of beer while learning about open science principles. Similarly, the BioDIGS project, part of the Genomic Data Science Community Network (GDSCN)^49^, empowers scientists from underserved institutions across the United States to investigate the microbial life in their local environments (https://biodigs.org/). Additionally, a non-profit organisation, SPUN (https://www.spun.earth/), leverages Microbiology Galaxy Lab to map mycorrhizal fungal communities globally. These diverse use cases highlight the Microbiology Galaxy Lab’s role in advancing microbial research, fostering collaboration, and making complex analyses accessible to researchers and citizen scientists.

Microbiome functional annotation seeks to define the role of constituent genes, often by linking genetic potential (metagenomics) with real-time activity (metatranscriptomics, metaproteomics). However, accurate functional assignment remains a major challenge. Workflows are crucial for this task, providing standardised pipelines that automate gene prediction by searching against multiple reference databases to build comprehensive functional profiles. Through annotations, trait discovery allows researchers to pinpoint novel enzymes and metabolic pathways from the microbiome with direct applications in biotechnology^50^.

The Microbiology Galaxy Lab provides workflows and tools to perform functional annotation and discover traits by utilising multi-omics data. A compelling example of this is the work by Péguilhan et al. 2025^29^ who combined metagenomics and metatranscriptomic data to reveal microbes functioning in different atmosphere compartments (**Fig 4B**). To overcome the high sample variability, they constructed an annotated gene catalogue from assembled metagenomes. This enabled a precise differential expression analysis, revealing key metabolic and stress responses triggered by the cloud water environment with implications for atmospheric chemistry and surface ecosystems likely to receive precipitation. A study of the North Atlantic Ocean microbiome^51^ underscored the importance of robust and reproducible workflows in environmental metaproteomics for accurate taxonomic and functional analyses of mass spectrometry data. In this context, the resources in the Microbiology Galaxy Lab contribute by providing clear guidelines and workflows for proteomics. Interlaced workflows, Schiml et al. 2023^52^ successfully demonstrated a comprehensive assessment of a community’s functional potential through the seamless multi-omic integration of metagenomic, metatranscriptomic, and metaproteomic data (**Fig 4C**). Their sophisticated strategy involved first recovering high-quality metagenome-assembled genomes (MAGs) and then using the predicted genes to construct custom databases for the subsequent protein and mRNA identification. Finally, metabolomics can complement these analyses by reflecting real-time cell function. The platform ecosystem is fully equipped to support such integrated analyses^53–55^.

### A Microbiology-focused Galaxy Lab for FAIR and reproducible analyses

The Microbiology Galaxy Lab is a Galaxy Lab. A Galaxy Lab is an extension to an existing Galaxy server that provides a custom landing page and a tailored toolbox when the server is accessed through a subdomain. This approach allows users to access domain-specific tools, workflows, and resources within the Galaxy environment^18^, i.e., a user-friendly interface for data analysis without an end user’s need for programming expertise. The Microbiology Galaxy Lab offers a curated interface for microbiology research, enabling users to run individual tools or build complex workflows, with full provenance tracking to ensure reproducibility. It is hosted on public Galaxy servers (microbiology.usegalaxy.org, .eu,.org<u>.au</u>, <u>.fr</u>), providing free access to storage and computational resources up to server-specific limits. This setup empowers users worldwide to perform sophisticated analyses regardless of local infrastructure. Users can upload data locally, via bulk data transfer or commercial clouds, or fetch it automatically from sources like UCSC Genome Browser database^56^, NCBI Sequence Read Archive (SRA)^57^, EMBL-EBI European Nucleotide Archive (ENA)^58^, and MGnify^59^. Automatic datasets retrieval and reuse, paired with Galaxy’s capability to export workflows as RO-Crate^60^ packages, ensures analysis reproducibility and machine-readability in line with FAIR guidelines. The Microbiology Galaxy Lab facilitates FAIR and reproducible microbial data analyses (**Table 1**).

**Table 1:** Support for FAIR and reproducible microbial data analyses in the Microbiology Galaxy Lab.

| FAIR Principles |  | Implementation in Microbiology Galaxy Lab |
| --- | --- | --- |
| Findable | Metadata and Tagging | Galaxy allows users to annotate datasets, workflows, and analyses with rich metadata (e.g., tags, descriptions, and ontologies like EDAM), |
|  | Persistent Identifiers (PIDs) | Galaxy supports the use of DOIs (Digital Object Identifiers) for workflows, ensuring they can be uniquely identified and cited. |
|  | Public Repositories | Workflows and tutorials published on platforms like WorkflowHub, Dockstore, and ELIXIR TeSS are indexed and searchable, increasing their visibility. |
| Accessible | Open Access | Galaxy is open-source and freely available, allowing anyone to install and use it. Public Galaxy servers (e.g., usegalaxy.org, usegalaxy.eu) provide free access to computational resources for researchers worldwide. |
|  | Data Sharing | Users can share datasets, workflows, and histories via shareable links or by publishing them to public repositories. Access permissions can be finely controlled (e.g., private, shared with collaborators, or fully public). |
|  | Cloud and Containerisation | Galaxy supports deployment in cloud environments (e.g., Galaxy on Kubernetes, CloudLaunch) and containerisation (e.g., Docker, Singularity), making it accessible across diverse infrastructures. |
| Interoperable | Adherence to community standards | Galaxy is an ELIXIR Recommended Interoperability Resource (RIR), underscoring its commitment to interoperability and community standards. Galaxy's commitment to interoperability is further demonstrated through its active implementation and support of Global Alliance for Genomics and Health (GA4GH) standards and APIs |
|  | Standardised Formats | Galaxy supports a wide range of standard data formats (e.g., FASTQ, BAM, VCF, Tabular) and encourages the use of community standards (e.g., GA4GH for genomics). |
|  | APIs and Interface | Galaxy provides REST APIs (OpenAPI annotated) and integrates with GA4GH APIs, including: (i) Data Repository Service (DRS) API as both a producer and consumer, facilitating easy exchange of genomic data with other DRS-compliant systems; (ii) Beacon API, enabling researchers to discover and query genomic variants across datasets; (iii) Task Execution Service (TES) API leveraged by Galaxy's Pulsar for large-scale genomic analyses, ensuring standardized and interoperable task execution; (iv) Tool Registry Service (TRS) API to access and share workflows via platforms like Dockstore and WorkflowHub, making it easy for users to find, import, and run community-developed |
|  |  | workflows. |
|  | Integration with Other Tools and Platforms | Galaxy supports integration with popular data science environments such as Jupyter Notebooks, RStudio, and ELIXIR services, enabling seamless data exchange and interoperability with external systems. |
|  | Workflow Standard | Galaxy workflows can be exported in RO-Crate format and are interoperable with other workflow engines. Galaxy also integrates with Dockstore and WorkflowHub, allowing users to search, import, and publish workflows directly from the Galaxy interface. |
| Reusable | Reproducibility | Galaxy captures the entire computational environment (tools, versions, parameters) for each analysis, ensuring that workflows can be rerun with identical results. This is critical for reproducibility. |
|  | Versioning | Workflows and tools in Galaxy are versioned, allowing users to track changes and revert to previous versions if needed. |
|  | Documentation | Galaxy encourages detailed documentation for workflows and tools, including step-by-step guides, a best practice wizard, tutorials via the Galaxy Training Network. |
|  | Community and Collaboration | The Galaxy community actively shares best practices, workflows, and tools, fostering reuse and collaboration. Platforms like Galaxy Europe and Galaxy Australia provide additional support and resources. |

### Resources backed by comprehensive training support

Microbiology Galaxy Lab facilitates users to execute complex analyses through precise self-guided training, providing understanding, learning, and enhancing data interpretation. The Lab offers 39 tutorials, 17 videos (16 hours), and 4 structured learning pathways for microbial analyses hosted on the Galaxy Training Network (GTN)^61,62^ (**Extended Data Fig. 4, Supplementary Table 6**). The GTN is an open, community-driven resource providing high-quality training materials and hands-on tutorials for learning data analysis and workflow usage within Galaxy. These resources cover various topics, from basic sequence analysis to advanced metaproteomics or metagenomics assembly (**Extended Data Fig. 4**), with regular updates ensuring alignment with the latest tools and methodologies. A steady cumulative increase in GTN tutorials with a microbial focus has occurred since 2016 with current growth of available materials ongoing (**Fig. 2A**). Integrating these materials into the Microbiology Galaxy Lab enhances user accessibility, experience, and confidence in conducting complex microbiology analyses.

Beyond self-guided resources, training events are critical in skill development. The microGalaxy community, behind the Microbiology Galaxy Lab, actively supports and leads diverse training efforts. Over the past five years, the microGalaxy community has conducted 30+ training events (**Supplementary Table 7**), often supported by the Galaxy Training Infrastructure as a Service (TIaaS) framework^63^, providing dedicated computing resources for efficient data processing during training sessions. A notable example is the 2021 *Analysis of Functions Expressed by Microbiomes* workshop (https://galaxyproject.org/events/2021-11-microbiomes/home/), co-hosted by CSIR-IMTech (Chandigarh, India) and Galaxy-P team (MN, USA), which introduced and trained 37 participants in microbiome functional analysis available while encouraging collaboration and knowledge sharing to promote microbial research impact.

Microbiology Galaxy Lab contains the knowledge and expertise from the Galaxy community and supports large-scale global initiatives like the Galaxy Training Academy (formerly Galaxy Smörgåsbord). Since 2021, this annual event has attracted over 10,000 participants, offering remote, asynchronous, video-based training with community support on the cloud-based communication platform, Slack (https://slack.com/). In 2024, the microGalaxy community offered two dedicated tracks in the annual Training Academy event: *Microbiome* and *Bacterial Genomics*, where 66.7% of 3,000+ registrants selected at least one of these tracks. In 2025, the *Microbiome* track was offered again, attracting 56.3% of a similarly large registrant base. The same model has been used for specialised trainings, such as *Mycobacterium tuberculosis* genomic analysis (https://training.galaxyproject.org/training-material/events/2024-06-10-mtb-ngs.html), reaching broad participation from countries in the Global South, where tuberculosis is a major health concern. These global events establish a strong foundation for correct workflow implementation, tool selection, and parameter tuning in the platform.

### Robust support and community engagement

Beyond training, Microbiology Galaxy Lab users benefit from strong support via the Galaxy Help forum (https://help.galaxyproject.org/) and a dedicated channel (https://matrix.to/#/#galaxyproject_microGalaxy:gitter.im) on the open communication network Matrix. The microGalaxy community, launched in 2021, fosters a collaborative atmosphere through regular meetings, working groups, and events. Hackathons, workshops, and meetings drive the new workflows development, tool improvements, and knowledge sharing. The microGalaxy community has led two ELIXIR BioHackathon projects^64^ and a dedicated microGalaxy hackathon^65^ to build the Galaxy Codex, a repository for Galaxy tools, workflows, and training metadata^66^. These efforts enhanced tool and tutorial annotations using the EDAM Ontology^67^ (see Methods). A hackathon in 2025 integrating workflows into the Intergalactic Workflow Commission (IWC) (https://github.com/galaxyproject/iwc) further expanded platform capabilities.

## Discussion

The Microbiology Galaxy Lab, supported by the microGalaxy community, provides a comprehensive, freely available (microbiology.usegalaxy.org, .eu, .org.au, .fr) platform for bringing microbial computational analysis to users. To facilitate advanced microbial research, the Microbiology Galaxy Lab integrates 318 analytical tool suites and 117 computational workflows with access to both public and local reference databases. This platform is supported by a rich training resources repository, including 39 tutorials and 17 videos, designed to support its widespread adoption and frequent utilisation. As an open-source solution within the Galaxy ecosystem, the platform provides extensive capabilities for microbial data analysis. It incorporates a workflow manager, an accessible interface, a comprehensive suite of curated analytical and visualization tools, and significant computational resources, thereby empowering researchers to perform complex analyses that might exceed their individual expertise or local computational capacity.

Over the past five years, the tools and workflows hosted in the Microbiology Galaxy Lab have been used in over 25 million analyses, reflecting their widespread acceptance and global reach. The platform’s robust infrastructure enables analyses ranging from short-read metabarcoding or genome assembly and taxonomic classification to complex metaproteomics and AMR surveillance workflows. This extensive usage highlights the platform’s critical role in supporting both routine and advanced microbiological research, with a user base spanning clinical, academic, and environmental institutions.

A key feature of Microbiology Galaxy Lab is the ability to respond to the changing landscape of microbiological data-intensive research. As microbial research evolves, so must the resources for data analysis. The Microbiology Galaxy Lab can be adapted alongside these developments by expanding the platform’s capabilities, improving multi-omics integration, and addressing emerging challenges. Platform adaptability is achieved through regular contributions from the global Galaxy community, including tool version upgrades, new microbial tool submissions, and curation through the microGalaxy community of practitioners. Dedicated initiatives, such as community hackathons, workshops, and collaborations with infrastructure partners like ELIXIR and the Galaxy Training Network, ensure that the platform evolves in line with scientific and technical developments.

The Microbiology Galaxy Lab provides the ideal foundation for using complex workflows that directly address critical challenges in health and disease. Its major strength lies in integrating complex, multi-step bioinformatic processes into accessible, end-to-end pipelines, making the platform the perfect ground for crucial analyses like AMR surveillance. Instead of requiring users to master individual command-line tools, the platform hosts complete workflows, such as AMR gene identification implemented in ABRomics that seamlessly guide analysis from raw patient sequence data to final AMR gene identification. By including important steps like quality control, genome assembly, and plasmid screening accessible through a user-friendly interface, Microbiology Galaxy Lab lowers the barrier to entry, helping clinicians and public health scientists to perform vital analyses for diagnostics but also for further research without needing specialised bioinformatics expertise.

Additionally, ecosystem and biodiversity research, especially in monitoring microbial communities in soils, oceans, and other ecosystems, can benefit from integrating microbial and ecological data analyses. Combining the microGalaxy community with the Galaxy-Ecology community (https://ecology.usegalaxy.eu/) provided a guidance framework for best practices in microbial ecological data analysis^68^, promoting strong interdisciplinary research. These efforts are particularly valuable for long-term environmental monitoring projects, enabling researchers to track microbial shifts in diverse ecosystems such as cloud microbiomes, root-associated plant environments, and fungal populations like Trichoderma species in soil. By enabling microbial and environmental data types integration, the platform strengthens our ability to interpret microbial roles in ecosystem services such as carbon cycling, pollutant degradation, and climate resilience. Looking ahead, a key direction for environmental research is a proper meta-omics integration, in particular with metabolomics, to enable systems-level insights. Supporting such multi-omics analyses within the Microbiology Galaxy Lab would enable genome-scale metabolic modelling, allowing researchers to infer microbial functions, interactions, and metabolic fluxes in complex ecosystems. This integration will help researchers apply standardised and reproducible approaches to linking microbial insights with ecological ones. As a result, it will enhance our ability to study microbial diversity in response to global changes, improving our ecosystem health and resilience understanding.

One Health, one of the microbiology focuses, is highly represented in the Microbiology Galaxy as illustrated by several use cases (**Supplementary Table 3)**. The major workflows to study antimicrobial resistance genes in diverse samples and locations, identify plasmids in pathogens, detect virulence factors, and/or track dysbiosis among clinical and environmental communities are all present and available to any user. Additionally, the possibility of FAIR data management contributes to generating clinical or fundamental conclusions regarding pathogens and antibiotic resistance mechanisms spread.

To enhance collaborative and in-depth microbial community analysis, we integrated QIIME(2), allowing users without computational expertise to perform microbiome analyses via a user-friendly workflow manager and training materials. Other future integrations, including Anvi’o^69^ and Nextstrain^70^, will enable more in-depth microbial community analysis. Federated data analysis is another priority, pursued through collaborations with groups like EMBL-EBI and the MGnify team to integrate all MGnify pipelines into Microbiology Galaxy Lab. This strategy scales computational capabilities and, similar to ABRomics, incorporates workflows into Bioinformatics Resource Centers (BRCs) like BRC Analytics (https://brc-analytics.org/) to enable seamless omic analyses of pathogenic organisms. While building on existing capabilities for viral analyses like important pandemic viruses SARS-CoV-2^71^ and Mpox, the platform can additionally include archaea and eukaryotic microbes to fill a critical research gap. Dedicated tools inclusion for these domains, which are often obscured by dominant prokaryotic signals^72^, strengthens support for multi-omic and holo-omic approaches, thereby enabling a more comprehensive host-microbiome interactions understanding through integrated (meta)omics data.

As microbial data generation continues to accelerate, it creates a critical need for the efficient computational resources allocation. The Microbiology Galaxy Lab has a dynamic computational resources allocation based on the Galaxy’s Total Respective Vortex (https://github.com/galaxyproject/total-perspective-vortex). But predictive tools developments estimating resource requirements from input datasets are indispensable to ensure that users can run large-scale analyses on available workflows without overloading resources. Computational power distribution allows flexibility in workflow implementation and improves users’ experience.

By providing a comprehensive suite of workflows, tools, and training resources representing the full spectrum of diverse and complex microbiome analyses, the Microbiology Galaxy Lab marks a fundamental shift from a static repository to a dynamic, community-driven ecosystem. It connects these powerful technologies within an accessible and collaborative framework, removing technical barriers and encouraging researchers to pursue questions previously beyond their reach. As a central platform for discovery, the platform is set to not only accelerate research pace but to fundamentally deepen our collective microbial world understanding. The Microbiology Galaxy Lab is therefore emerging as a microbiology cornerstone, not only by serving hundreds of thousands of users but by setting a new standard for collaborative research platforms across all scientific disciplines.

## Materials and methods

### Galaxy framework and the Microbiology Galaxy Lab

The Galaxy framework^18^ is a robust and widely adopted platform enabling accessible and reproducible bioinformatics analyses. The Microbiology Galaxy Lab extends the Galaxy framework using a domain-specific interface called a Galaxy Lab. A Galaxy Lab, built and deployed with the Galaxy framework and Galaxy Labs Engine (https://labs.usegalaxy.org.au)^73^, provides users with a focused workspace retaining all the benefits of the Galaxy ecosystem, including accessibility, scalability, and reproducibility. With templated elements, each server can host a similar version of a Galaxy Lab, centralising community resources while allowing adaptations such as server-specific branding and support links.

The Microbiology Galaxy Lab is deployed as a dedicated subdomain on several public Galaxy servers (usegalaxy.eu, usegalaxy.org, usegalaxy.org.au, usegalaxy.fr). This availability provides users with streamlined access to its suite of tools and workflows for microbiological data analysis. Furthermore, it leverages the robust computational resources of these host infrastructures, thereby removing the need for local software installation or specialised hardware on the part of the end-user.

All resources supporting the Microbiology Galaxy Lab, including its customised interface, and lists of tools, workflows, and training materials, are curated and stored within the Galaxy CoDex GitHub repository (https://github.com/galaxyproject/galaxy_codex). The Galaxy CoDex is a centralised repository ensuring the versioning and documentation of the Microbiology Galaxy Lab components.

### Community-curated tools

Focused on microbial research, the Microbiology Galaxy Lab facilitates access to tool suites, workflows, and training materials essential for microbial data analysis. Available tools are supported by the microGalaxy community, an engaged group of practitioners experienced in microbial data analysis. Tools are sourced from the Galaxy ToolShed^74^ (https://toolshed.g2.bx.psu.edu), a central repository hosting tools wrapped and made available for deployment on the Galaxy platform, facilitating their straightforward access and installation on various Galaxy servers, similar to an App Store. The tool wrappers provide the integration layer between the software or tool command-line and the Galaxy platform, defining inputs/outputs (including their formats) and parameters. Furthermore, they are developed and maintained by the global Galaxy community and groups such as the Intergalactic Utilities Commission (IUC) (https://galaxyproject.org/iuc/), with their source code managed in open-source, publicly available GitHub repositories (https://github.com/galaxyproject/planemo-monitor). Galaxy tool wrapper development, testing, and validation are conducted using Planemo^75^, a command-line Software Development Kit (SDK). The use of Planemo promotes the creation of consistent, high-quality tool wrappers for the Galaxy platform. Following development and community approval, these tools are deposited in the Galaxy ToolShed.

Any tool dependencies are resolved using packages or containers available through Bioconda^76^ or Biocontainers^77^. When the dependencies are updated, tools maintained by the IUC or other community tool repositories undergo a semi-automated updating process. Once approved by a community member, the updated tool becomes publicly available on ToolShed and is automatically updated on the major Galaxy servers where they are installed. This process ensures that users have access to the latest versions of tools. At the same time, legacy versions are retained to support reproducibility and maintain standardised workflows, enabling scientists and clinical centers to repeat analyses even as tools evolve. Tools in the Galaxy ToolShed are organised into individual entries, each representing a single command, or grouped into tool suites, which are sets of related software commands.

In the Microbiology Galaxy Lab, these tool suites are carefully selected based on their relevance to microbial research, their widespread use within the community, and their adherence to Galaxy’s development best practices. The selection process is guided by input from experts within the microGalaxy community, ensuring the inclusion of robust, well-maintained tools that are essential for a variety of workflows, from genome assembly to complex tasks like antimicrobial resistance analysis. Curation is managed through the Galaxy CoDex GitHub repository using a semi-automated process: weekly automatic updates add new, potentially interesting tools to a review list, and community members regularly evaluate this list. Approved tools are then added to the Community-curated Tools list, maintaining an up-to-date and authoritative collection.

### Community-curated workflows

Workflows included in the Microbiology Galaxy Lab are selected based on their relevance to microbial research, adherence to Galaxy best practices, compatibility with training material or peer-reviewed studies, and community adoption metrics such as usage and demand. The Microbiology Galaxy Lab workflows exist at four levels of development, providing varying degrees of accessibility, adherence to best practices, and maintenance: (i) Publicly shared workflow: These workflows, tagged with #microGalaxy, require minimal effort to create. They are readily accessible to the research community, available across public Galaxy instances, and offer a starting point for users seeking to conduct microbial data analysis; (ii) Workflows in WorkflowHub^24^: Workflows published in WorkflowHub, a workflow registry, are annotated with metadata such as creator information and licensing. These workflows may not include test data, but they offer documented and structured analysis workflows for microbial research; (iii) Workflows in WorkflowHub related to tutorials: workflows supported by tutorials on the Galaxy Training Network (GTN)^61,62^. These workflows adhere to Galaxy’s best practice, i.e., including clear annotations, proper input and output parameter definitions, formal documentation (such as creator information and licensing), and test data; and (iv) IWC-Validated Workflows on WorkflowHub and Dockstore^25^: A subset of workflows available through WorkflowHub has been further curated through the Intergalactic Workflow Commission (IWC) (https://github.com/galaxyproject/iwc) to adhere to Galaxy’s best practices, including test data, and support state-of-the-art analyses. Stored in the IWC GitHub repository, they are supported by a semi-automated updating system. When a Galaxy tool used in the workflow is updated, a continuous integration pipeline tests the workflow with the updated tool version. Once a community member approves, the updated workflow becomes publicly available on WorkflowHub, Dockstore, and the major Galaxy instances. This process ensures that microGalaxy community workflows remain up-to-date and reproducible, making them an invaluable resource for microbial research. Galaxy users can easily import workflows from WorkflowHub or Dockstore directly into their Galaxy server, enabling seamless integration into their analysis pipelines.

Similar to the tools, the curation process for the four distinct workflow types is handled through the Galaxy CoDex GitHub repository via a semi-automated system. This process involves weekly automatic updates that introduce new, potentially valuable workflows to a review list. Community members then regularly assess this list, and those workflows that are approved are added to the Community-curated Workflows list, ensuring the collection remains current and authoritative.

### Training materials

Training is critical to equipping users with the knowledge needed to fully leverage the platform’s capabilities. The global Galaxy community has developed a comprehensive suite of over 400 tutorials, which are available on the GTN. The Microbiology Galaxy Lab curates tutorials by tagging microbial-focused ones with the keyword “microGalaxy” (https://training.galaxyproject.org/training-material/search2?query=microgalaxy). These tutorials cover various topics, from 16S rDNA gene amplicon analysis (https://gxy.io/GTN:T00441) to more complex analyses like metagenomics (https://gxy.io/GTN:T00385) or metaproteomics (https://gxy.io/GTN:T00221). The GTN tutorials include step-by-step instructions and test data. They are designed to support learners of varying expertise levels, from beginners to experienced researchers. Many tutorials are also supported by captioned video recordings hosted on YouTube to support flexible learning for a broader audience.

The Microbiology Galaxy Lab tutorials are continually updated to incorporate state-of-the-art tools, workflows, and methodologies, ensuring users can access current information. Many tutorials are structured into learning pathways, providing a step-by-step progression of topics, guiding learners through increasingly complex analyses. These learning pathways are designed to build foundational knowledge while advancing skills for handling cutting-edge microbial data. In addition to the available tutorials, Microbiology Galaxy Lab promotes a range of training events, such as workshops, hackathons, and online seminars focusing on current topics related to microbiology. These events allow users to learn, engage directly with bioinformatics experts, share experiences, and deepen their understanding of microbial computational analyses.

### Resource aggregation and annotation

Resource annotation and ontologies are essential for ensuring consistency and improving discoverability across the Microbiology Galaxy Lab. To achieve this, Microbiology Galaxy Lab employs the EDAM ontology^67^, a structured vocabulary for bioinformatics concepts, to categorise tools, workflows, and training materials. This ontology-based approach ensures consistent descriptions across the platform, streamlining resource discovery and enabling users to quickly find resources tailored to their research needs.

All Microbiology Galaxy Lab resources are aggregated into the Galaxy CoDex (https://github.com/galaxyproject/galaxy_codex), a comprehensive catalog that integrates metadata from the Galaxy ecosystem, along with the international registries for bioinformatic tools, bio.tools^78,79^ (https://bio.tools/), and workflows, WorkflowHub^24^ (https://workflowhub.org/). The CoDex enables the creation of lists and widgets that can be embedded into websites, offering seamless access to up-to-date tools, workflows, and training materials.

During a resource aggregation phase, missing tool annotations were identified, particularly for tools lacking EDAM ontology terms or links to software registries such as bio.tools. To address this, an annotation process was employed in Microbiology Galaxy Lab, supported by the microGalaxy community, including dedicated hackathons^64,65^. These efforts resulted in over 315 tool suites being annotated and more than 35 tutorials being enriched with relevant EDAM terms. This process improved the categorisation and discoverability of resources within the Microbiology Galaxy Lab and provides a model for other scientific communities to follow in organising their resources.

### Survey and use cases

A survey was conducted between March and September 2023 to assess the needs, preferences, and challenges of the microbial research community. The survey aimed to gather insights into how researchers interact with the Galaxy platform for microbial research and identified areas for improvement. The survey was developed by the microGalaxy community and was distributed online via the Galaxy Project website, mailing lists, and social media channels. It consisted of 16 questions (**Supplementary Document 1**) that covered several categories to understand end users, including (i) research focus and community demographics, (ii) tools and workflows, (iii) training and support, and (iv) future developments. The survey included multiple-choice, Likert scale, and open-ended questions to allow the collection of both quantitative and qualitative information^80^.

Participation was voluntary, and participants were invited to respond after an introduction to the survey goals. A total of 130 respondents (**Supplementary Table 1**) participated in the survey, representing a diverse range of geographical locations, institutions, and research domains. Quantitative responses were analysed using descriptive statistics to summarise trends and identify prevalent themes. Qualitative responses to open-ended questions were coded and analysed thematically to extract insights into specific challenges and recommendations.

Survey participants who indicated a willingness to be contacted were sent a follow-up invitation, including a structured document (**Supplementary Document 2**) to collect detailed information about their use cases. This document included (i) research objectives and questions, (ii) methods employed, including experimental techniques, data generation approaches, and analysis pipelines, and (iii) tools and workflows used within and outside Galaxy. The 21 use cases were collected and anonymised (**Supplementary Table 3**). The compelling use cases were elaborated upon in the main text of this study, and all authors were invited to contribute to the manuscript preparation.

### Citation extraction and annotation

To analyse the impact of Galaxy on microbial research, citations of the major Galaxy papers were extracted as described in **Extended Data Fig. 5** using Python Programming Language (v3.8.19, RRID:SCR_008394) within a Jupyter Notebook (v1.0.0, RRID:SCR_018315). First, the 8 major publications of the Galaxy Project were extracted from the Galaxy Project’s Google Scholar profile (https://scholar.google.com/citations?hl=en&user=3tSiRGoAAAAJ) using the scholarly package (v 1.7.11)^81^. Second, citations of these publications were then retrieved on Semantic Scholar^82^ via its Application Programming Interface (API) using requests (v2.32.3). The collected data included the publication years, titles, and abstracts.

Citations of the 8 major publications of the Galaxy project were annotated as microbial-related if their titles or abstracts contained at least one of the 25 predefined keywords relevant to microbial research. These microbial-related citations were then associated with topics; “Ecosystem Dynamics & Biodiversity” and/or “Health & Disease” if titles or abstracts contain defined keywords for each topic. If neither title nor abstracts contain the respective keywords, both “Ecosystem Dynamics & Biodiversity” and “Health & Disease” topics are assigned. Parallelly microbial-related citation was further categorised into other three dimensions given keywords in their titles and abstracts to enable a detailed analysis of the research themes addressed in the citing papers: (i) Targeted Organisms (e.g., Bacteria, Virus), (ii) Technical Targets (e.g., Isolate, Community (taxonomy) profiling), and (iii) Methods (e.g., Metabarcoding, (Meta)genomics(Meta)transcriptomics).

### Data visualisation

To analyse and visualise summary data presented in this study, a set of scripts in RMarkdown (https://github.com/rstudio/rmarkdown) was created. These scripts streamlined the generation of figures and statistical analyses, allowing for reproducibility and updates as new data became available. Each script ingests tables generated by Galaxy CoDex, representing the Microbiology Galaxy Lab resources and containing metadata and EDAM annotations for tools, training materials, and workflows. This setup enables the calculation of key metrics such as total counts, distribution across EDAM terms, and user feedback statistics.

All analyses were run on R (v4.3.1, RRID:SCR_001905) with the use of *data.table* (v1.14.8, [RRID:SCR_026117](https://rrid.site/data/record/nlx_144509-1/SCR_001905/resolver?q=SCR_001905&l=SCR_001905&i=rrid:scr_001905)) and *tidyr* (v1.1.3, RRID:SCR_017102) libraries for data manipulation, and the *stringr* (v1.5.0, RRID:SCR_022813) library for text processing. Visualisations were primarily created with *ggplot2* (v3.4.4, RRID:SCR_014601), while *ggrepel* (v0.9.3, RRID:SCR_017393), *ggtext* (v0.1.2, RRID:SCR_026470), and *ggh4x* (v0.2.5) were used for labelling, rich text formatting, and facet customisation, respectively. The library *patchwork* (v1.3.2, RRID:SCR_024826), was used to compose plots into the same graphic, and the libraries, *colorspace,* and *paletteer* (v2.1-0) ensured accessible colour schemes, where *packcircles* (v0.3.5) handled circle-packing layouts, and *shadowtext* (v0.1.2) improved label readability. Additionally, *extrafont* (v0.18) was used to manage custom fonts.

## Code availability

The code used to conduct the data analyses and generate the figures presented in this paper can be found on a dedicated GitHub repository (https://github.com/usegalaxy-eu/microbiology_galaxy_lab_paper_2025). The Microbiology Galaxy Lab resources, including sources, tools, workflows, and related code, are integrated into the Galaxy CoDex GitHub repository (https://github.com/galaxyproject/galaxy_codex).

## Data availability

All supplementary tables associated with this study are available in the dedicated GitHub repository (https://github.com/usegalaxy-eu/microbiology_galaxy_lab_paper_2025). To ensure reproducibility and long-term accessibility, a versioned release of the dataset was created on Zenodo^83^ on March 26th, 2025. Any updates to the repository will be tracked, but the version archived on Zenodo corresponds to the data used in this study.

## Acknowledgments

The microGalaxy community efforts and the Microbiology Galaxy Lab are made possible by a growing Galaxy community of worldwide users, developers, system administrators, and educators. The authors acknowledge the support of the entire European Galaxy Team. We are grateful to de.NBI-Cloud (German National Bioinformatic Research Infrastructure) and the UFR-RZ for hosting https://usegalaxy.eu, to the [IFB (Institut Francais de Bioinformatique)](https://www.france-bioinformatique.fr/en/home/) NNCR Cluster Task force for hosting https://usegalaxy.fr, François Enault (University of Clermont Auvergne), Alexander von Humboldt Foundation.

Generative AI tools (ChatGPT 4o; July 2025) were used to revise and improve the clarity of the manuscript text. None of the manuscript content was generated entirely by AI.

## Extended data figures

**Extended Data Fig. 1:**
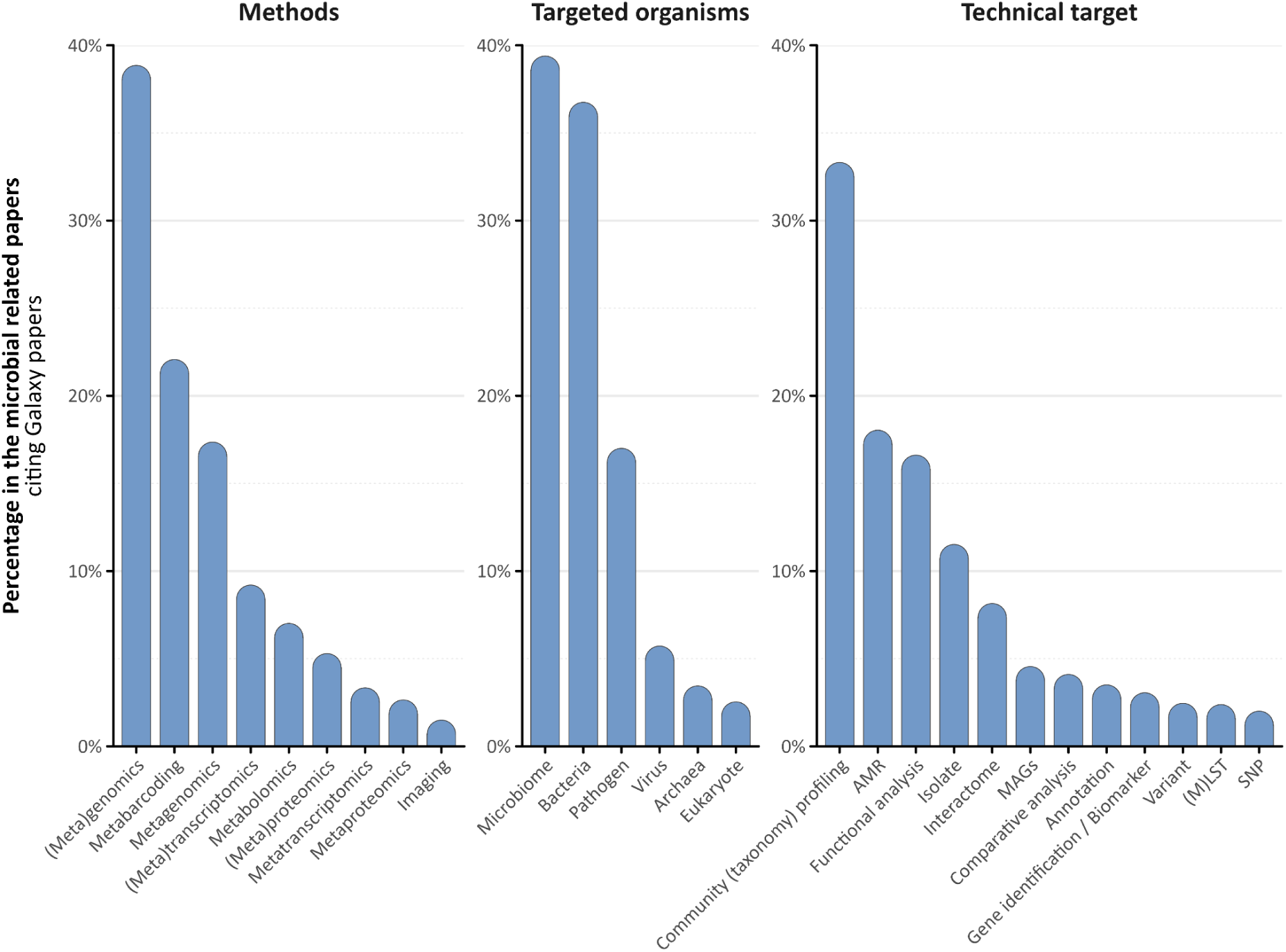
Microbial Research Topics in Galaxy Publications. Breakdown of microbial-focused citations by Targeted Organisms, Technical Targets, and Methods. Categories were annotated based on predefined keywords found in the title or abstract of each publication.

**Extended Data Fig. 2:**
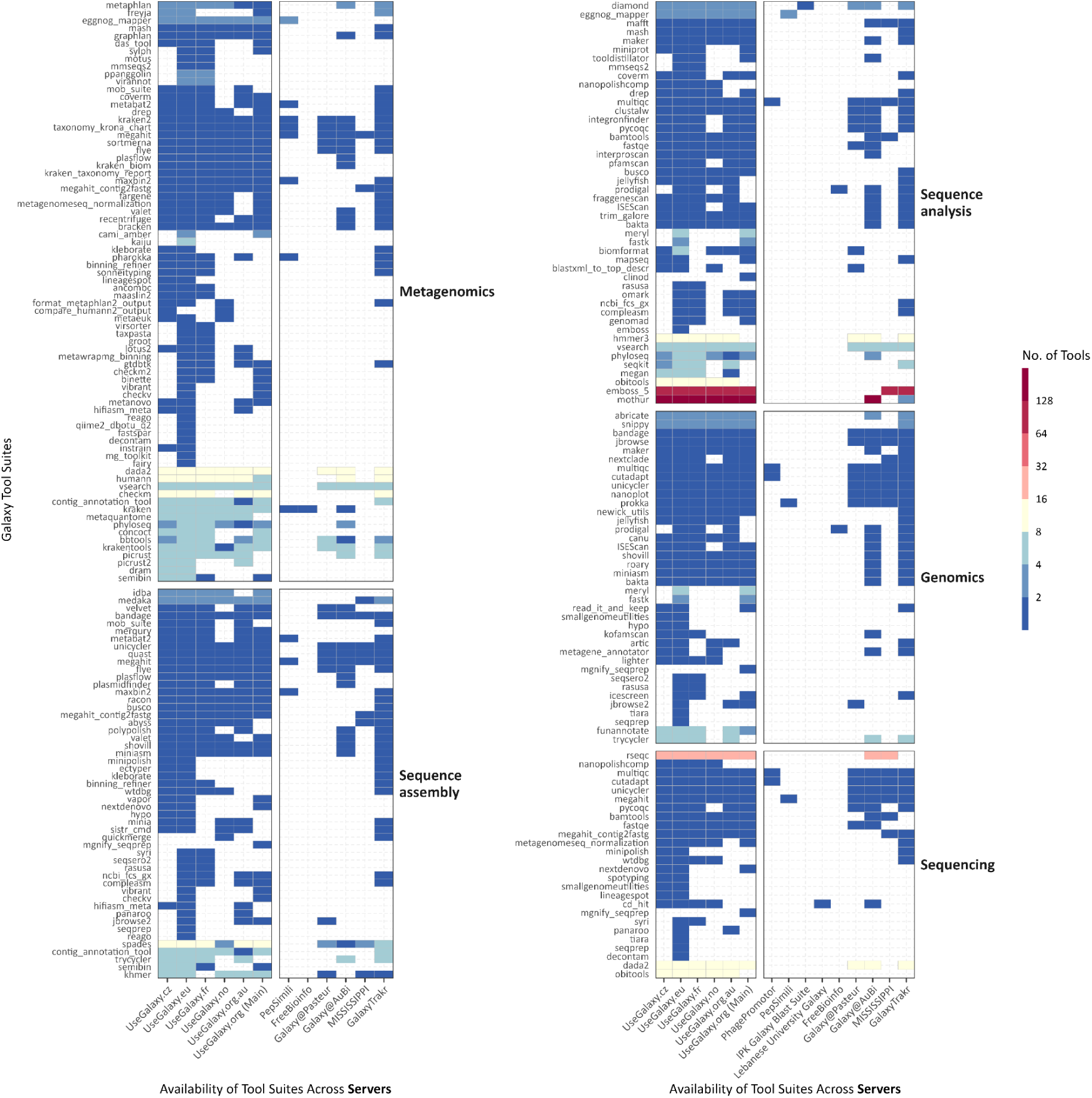
Availability of microbiology-related tool suites within the Galaxy ecosystem. Heatmap illustrating the availability of microbiology-related tool suites (x-axis) across various Galaxy servers (y-axis), grouped by EDAM topics. Tool suites may appear in multiple groups as they can be annotated with several topics. A logarithmic scale is applied for improved visualisation.

**Extended Data Fig. 3:**
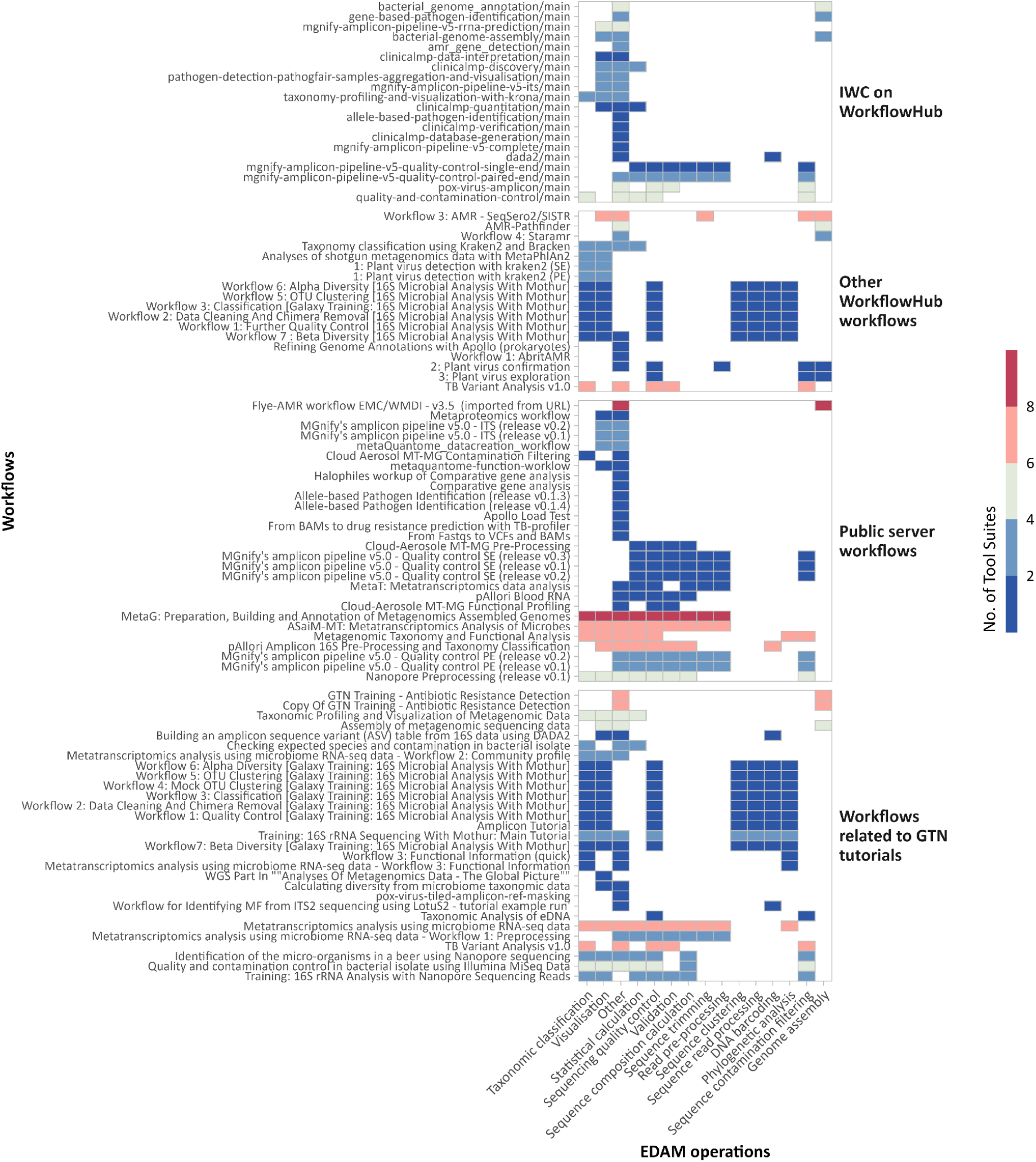
Usage of microbiology-related tool suites across workflows. Heatmap illustrating the presence of microbiology-related tool suites within the available microbiology-related workflows, grouped by four levels of development. The tool suites are organised based on their corresponding EDAM operations, highlighting the breadth of tools utilised for different workflows.

**Extended Data Fig. 4:**
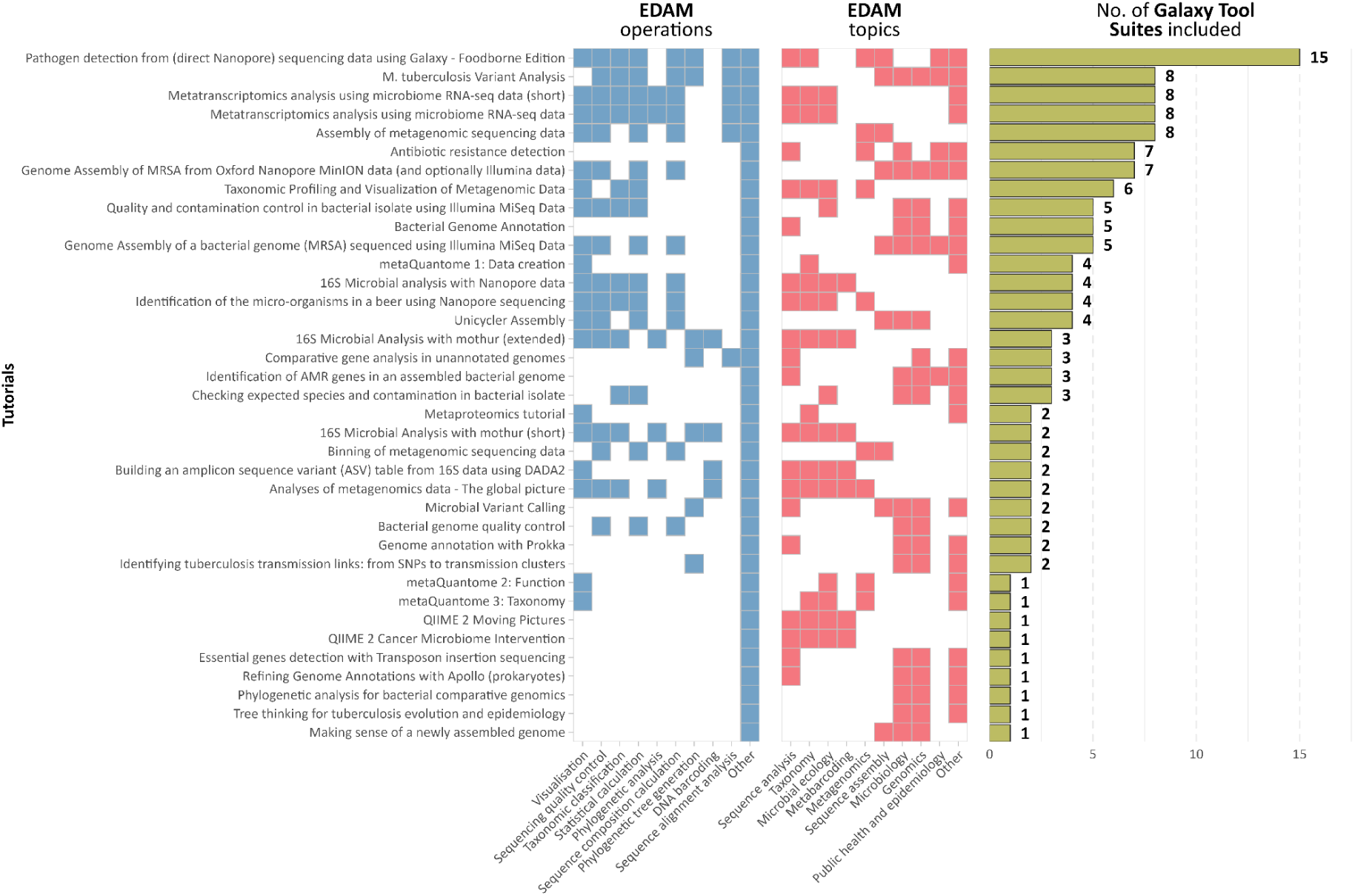
Usage of microbiology-related tool suites across training materials. Heatmap illustrating the presence of microbiology-related tool suites within the available microbiology-related training materials, grouped by EDAM operations (blue), EDAM topics (red), and number of Galaxy tool suites used in the tutorials (green), highlighting the breadth of tools utilised for different training contexts.

**Extended Data Fig. 5:**
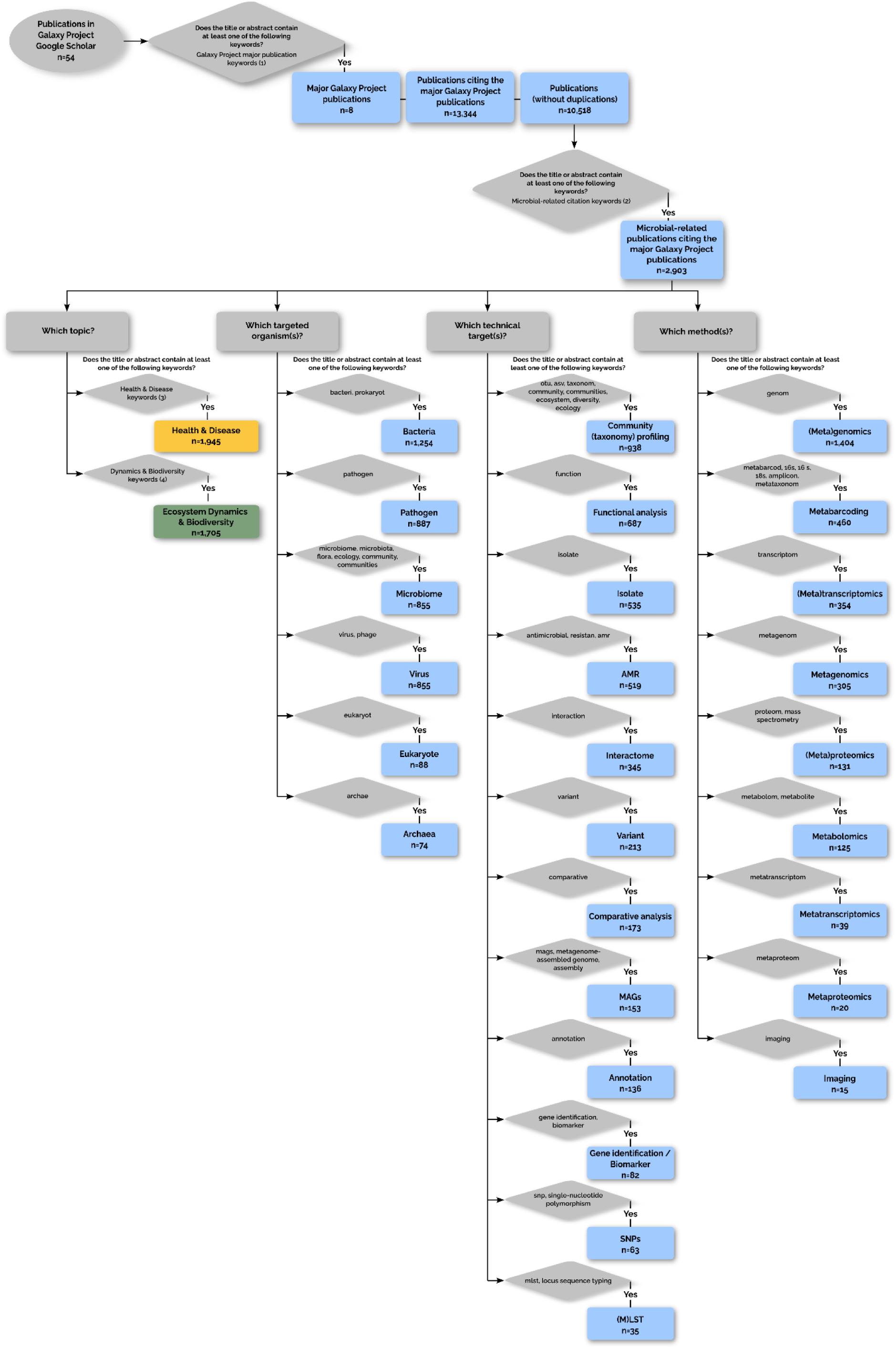
Flowchart illustrating the process to select and annotate the microbial-related publications citing the Galaxy Project major publication. After choosing the major Galaxy Project publications by filtering publications present in Galaxy Project Google Scholar based on predefined keywords (List 1 below) found in titles or abstracts, the publications citing them are extracted and duplications are removed. The unique publications are first screened for microbial relevance based on 25 predefined keywords (List 2 below) found in titles or abstracts. Microbial-related citations were classified into topic(s), targeted organism(s), technical target(s), and method(s) based on the presence or absence of specific thematic keywords, numbered or mentioned directly on Figure. Lists of keywords: (1) Galay Project major citations keywords: ‘platform’, ‘a comprehensive approach’, ‘a web-based genome analysis tool for experimentalists’. (2) Microbial-related citations keywords: ‘bacteri’, ‘prokaryot’, ‘microb’, ‘pathogen’, ‘virus’, ‘phage’, ‘archae’, ‘flora’, ‘microecology’, ‘microorganism’, ‘micro-organism’, ‘microbiome’, ‘microbiota’, ‘metabarcod’, ‘16s’, ‘18s’, ‘amplicon’, ‘metataxonom’, ‘metagenom’, ‘metatranscriptom’, ‘metaproteom’, ‘multi-locus sequence typing’, ‘multilocus sequence typing’, ‘mlst’, ‘otu’. (3) Health & Disease keywords: ‘health’, ‘disease’, ‘pathogen’, ‘antibiotic’, ‘epidemio’, ‘clinic’, ‘syndrom’, ‘antimicrob’, ‘patient’, ‘cancer’, ‘covid’, ‘hiv’, ‘diagnostic’, ‘food safety’, ‘toxic’, ‘infest’, ‘infect’, ‘prebiotic’, ‘probiotic’, ‘obesity’, ‘immun’, ‘food safety’, ‘gut microbiota’, ‘dysbiosis’, ‘host-microbe’, ‘immunomodulation’, ‘inflammation’, ‘gut-brain axis’, ‘metabolic syndrome’, ‘FMT’, ‘SCFA’, ‘healthcare’, ‘virulence’, ‘drug’. (4) Dynamics & Biodiversity keywords: ‘environment’, ‘diversity’, ‘ecosystem’, ‘plant’, ‘corrosion’, ‘geograph’, ‘compos’, ‘fish’, ‘soil’, ‘biofilm’, ‘water’, ‘aquaculture’, ‘ecolog’, ‘ferment’, ‘ocean’, ‘marine’, ‘decomp’, ‘biofuel’, ‘permafrost’, ‘microbial mat’, ‘rhizosphere’, ‘grassland’, ‘biodegrada’, ‘biogeochem’, ‘succession’, ‘trophic’, ‘food web’, ‘community assembly’, ‘resilience’, ‘stability’, ‘niche’, ‘alpha diversity’, ‘beta diversity’, ‘symbiosis’, ‘competition’, ‘functional diversity’, ‘phylogen’, ‘river’, ‘crop’, ‘bird’, ‘cyanobacteri’, ‘auquatic’.

## References

1. Di Bella, J. M., Bao, Y., Gloor, G. B., Burton, J. P. & Reid, G. High throughput sequencing methods and analysis for microbiome research. J. Microbiol. Methods 95, 401–414 (2013).

2. Reuter, J. A., Spacek, D. V. & Snyder, M. P. High-Throughput Sequencing Technologies. Mol. Cell 58, 586–597 (2015).

3. Filardo, S., Di Pietro, M. & Sessa, R. Current progresses and challenges for microbiome research in human health: a perspective. Front. Cell. Infect. Microbiol. 14, (2024).

4. Goodswen, S. J. et al. Machine learning and applications in microbiology. FEMS Microbiol. Rev. 45, fuab015 (2021).

5. Berrios, D. C., Beheshti, A. & Costes, S. V. FAIRness and Usability for Open-access Omics Data Systems. AMIA Annu. Symp. Proc. AMIA Symp. 2018, 232–241 (2018).

6. Wilkinson, M. D. et al. The FAIR Guiding Principles for scientific data management and stewardship. Sci. Data 3, 160018 (2016).

7. Ladner, J. T., Grubaugh, N. D., Pybus, O. G. & Andersen, K. G. Precision epidemiology for infectious disease control. Nat. Med. 25, 206–211 (2019).

8. Köser, C. U. et al. Whole-Genome Sequencing for Rapid Susceptibility Testing of *M. tuberculosis*. N. Engl. J. Med. 369, 290–292 (2013).

9. Khan, M. M. et al. Multi-Omics Strategies Uncover Host–Pathogen Interactions. ACS Infect. Dis. 5, 493–505 (2019).

10. Deng, L., Yin, H., Tan, K. S. W. & Tsaousis, A. D. Editorial: Zoonotic diseases: epidemiology, multi-omics, and host-pathogen interactions. Front. Microbiol. 15, (2024).

11. Romanis, C. S., Timms, V. J., Nebauer, D. J., Crosbie, N. D. & Neilan, B. A. Microbiome analysis reveals *Microcystis* blooms endogenously seeded from benthos within wastewater maturation ponds. Appl. Environ. Microbiol. 90, (2024).

12. Shaffer, J. P. et al. Standardized multi-omics of Earth’s microbiomes reveals microbial and metabolite diversity. Nat. Microbiol. 7, 2128–2150 (2022).

13. Keneally, C. et al. Multi-omics reveal microbial succession and metabolomic adaptations to flood in a hypersaline coastal lagoon. Water Res. 280, 123511 (2025).

14. Prieto Riquelme, M. V., et al. Demonstrating a Comprehensive Wastewater-Based Surveillance Approach That Differentiates Globally Sourced Resistomes. Environ. Sci. Technol. 56, 14982–14993 (2022).

15. Spänig, S. et al. A multi-omics study on quantifying antimicrobial resistance in European freshwater lakes. Environ. Int. 157, 106821 (2021).

16. One Health High-Level Expert Panel (OHHLEP) et al. One Health: A new definition for a sustainable and healthy future. PLOS Pathog. 18, e1010537 (2022).

17. Straub, D. et al. Interpretations of Environmental Microbial Community Studies Are Biased by the Selected 16S rRNA (Gene) Amplicon Sequencing Pipeline. Front. Microbiol. 11, (2020).

18. The Galaxy Community. The Galaxy platform for accessible, reproducible, and collaborative data analyses: 2024 update. Nucleic Acids Res. 52, W83–W94 (2024).

19. Blomer, J., Buncic, P. & Fuhrmann, T. CernVM-FS: delivering scientific software to globally distributed computing resources. in Proceedings of the first international workshop on Network-aware data management 49–56 (Association for Computing Machinery, New York, NY, USA, 2011). doi:10.1145/2110217.2110225.

20. Breitwieser, F. P. & Salzberg, S. L. Pavian: interactive analysis of metagenomics data for microbiome studies and pathogen identification. Bioinformatics 36, 1303–1304 (2020).

21. Bik, H. M. & Inc, P. I. Phinch: An interactive, exploratory data visualization framework for –Omic datasets. 009944 Preprint at 10.1101/009944 (2014).

22. McMurdie, P. J. & Holmes, S. Shiny-phyloseq: Web application for interactive microbiome analysis with provenance tracking. Bioinformatics 31, 282–283 (2015).

23. Lewis, S., et al. Apollo: a sequence annotation editor. Genome Biol. 3, research0082.1 (2002).

24. Gustafsson, O. J. R. et al. WorkflowHub: a registry for computational workflows. Sci. Data 12, 837 (2025).

25. Yuen, D. et al. The Dockstore: enhancing a community platform for sharing reproducible and accessible computational protocols. Nucleic Acids Res. 49, W624–W632 (2021).

26. Di Tommaso, P. et al. Nextflow enables reproducible computational workflows. Nat. Biotechnol. 35, 316–319 (2017).

27. Mölder, F. et al. Sustainable data analysis with Snakemake. F1000Research 10, 33 (2021).

28. Arcari, G. et al. Genotypic Evolution of Klebsiella pneumoniae Sequence Type 512 during Ceftazidime/Avibactam, Meropenem/Vaborbactam, and Cefiderocol Treatment, Italy. Emerg. Infect. Dis. 29, 2266 (2023).

29. Péguilhan, R. et al. Clouds influence the functioning of airborne microorganisms. Biogeosciences 22, 1257–1275 (2025).

30. Schiml, V. C. et al. Integrative meta-omics in Galaxy and beyond. *Environ*. Microbiome 18, (2023).

31. Pyles, R. B. et al. The altered TBI fecal microbiome is stable and functionally distinct. Front. Mol. Neurosci. 17, 1341808 (2024).

32. Do, K., et al. A novel clinical metaproteomics workflow enables bioinformatic analysis of host-microbe dynamics in disease. mSphere 9, (2024).

33. Walsh, T. R., Gales, A. C., Laxminarayan, R. & Dodd, P. C. Antimicrobial Resistance: Addressing a Global Threat to Humanity. PLOS Med. 20, e1004264 (2023).

34. Strepis, N. et al. BenchAMRking: a Galaxy-based platform for illustrating the major issues associated with current antimicrobial resistance (AMR) gene prediction workflows. BMC Genomics 26, 27 (2025).

35. Sherry, N. L. et al. An ISO-certified genomics workflow for identification and surveillance of antimicrobial resistance. Nat. Commun. 14, 60 (2023).

36. Sharma, T., et al. Galaxy ASIST: A web-based platform for mapping and assessment of global standards of antimicrobial susceptibility: A case study in Acinetobacter baumannii genomes. Front. Microbiol. 13, 1041847 (2023).

37. Merdan, O. et al. Investigation of the Defective Growth Pattern and Multidrug Resistance in a Clinical Isolate of Candida glabrata Using Whole-Genome Sequencing and Computational Biology Applications. Microbiol. Spectr. 10, e00776–22 (2022).

38. Bihani, S. et al. Metaproteomic Analysis of Nasopharyngeal Swab Samples to Identify Microbial Peptides in COVID-19 Patients. J. Proteome Res. 22, 2608–2619 (2023).

39. Kruk, M. E. et al. An integrated metaproteomics workflow for studying host-microbe dynamics in bronchoalveolar lavage samples applied to cystic fibrosis disease. mSystems 9, e00929–23 (2024).

40. Nasr, E., Henger, A., Grüning, B., Zierep, P. & Batut, B. PathoGFAIR: a collection of FAIR and adaptable (meta)genomics workflows for (foodborne) pathogens detection and tracking. Preprint at 10.1101/2024.06.26.600753 (2024).

41. Cumbo, F., Truglia, S., Weitschek, E. & Blankenberg, D. Feature selection with vector-symbolic architectures: a case study on microbial profiles of shotgun metagenomic samples of colorectal cancer. Brief. Bioinform. 26, bbaf177 (2025).

42. Rousk, J. & Bengtson, P. Microbial regulation of global biogeochemical cycles. Front. Microbiol. 5, (2014).

43. Gougoulias, C., Clark, J. M. & Shaw, L. J. The role of soil microbes in the global carbon cycle: tracking the below-ground microbial processing of plant-derived carbon for manipulating carbon dynamics in agricultural systems. J. Sci. Food Agric. 94, 2362–2371 (2014).

44. Bharti, R. & Grimm, D. G. Current challenges and best-practice protocols for microbiome analysis. Brief. Bioinform. 22, 178–193 (2021).

45. Hu, B. et al. Challenges in Bioinformatics Workflows for Processing Microbiome Omics Data at Scale. Front. Bioinforma. 1, 826370 (2022).

46. Rosolen, R. R. et al. Whole-genome sequencing and comparative genomic analysis of potential biotechnological strains of Trichoderma harzianum, Trichoderma atroviride, and Trichoderma reesei. Mol. Genet. Genomics 298, 735–754 (2023).

47. Kumaishi, K. et al. High throughput method of 16S rRNA gene sequencing library preparation for plant root microbial community profiling. Sci. Rep. 12, (2022).

48. Paul Zierep, EMBL’s European Bioinformatics Institute & Rand Zoabi. github.com/iwc-workflows/mgnify-amplicon-pipeline-v5-complete/main. Zenodo 10.5281/ZENODO.15078460 (2025).

49. Network, T. G. D. S. C. et al. Diversifying the genomic data science research community. Genome Res. 32, 1231–1241 (2022).

50. Prayogo, F. A. et al. Metagenomic applications in exploration and development of novel enzymes from nature: a review. J. Genet. Eng. Biotechnol. 18, 39 (2020).

51. Saito, M. A. et al. Results from a multi-laboratory ocean metaproteomic intercomparison: effects of LC-MS acquisition and data analysis procedures. Biogeosciences 21, 4889–4908 (2024).

52. Schiml, V. C. et al. Integrative meta-omics in Galaxy and beyond. *Environ*. Microbiome 18, 56 (2023).

53. Peters, K. et al. PhenoMeNal: processing and analysis of metabolomics data in the cloud. GigaScience 8, giy149 (2019).

54. Giacomoni, F. et al. Workflow4Metabolomics: a collaborative research infrastructure for computational metabolomics. Bioinforma. Oxf. Engl. 31, 1493–1495 (2015).

55. Davidson, R. L., Weber, R. J. M., Liu, H., Sharma-Oates, A. & Viant, M. R. Galaxy-M: a Galaxy workflow for processing and analyzing direct infusion and liquid chromatography mass spectrometry-based metabolomics data. GigaScience 5, 10 (2016).

56. Nassar, L. R. et al. The UCSC Genome Browser database: 2023 update. Nucleic Acids Res. 51, D1188–D1195 (2023).

57. Sayers, E. W. et al. Database resources of the national center for biotechnology information. Nucleic Acids Res. 50, D20–D26 (2022).

58. Burgin, J. et al. The European Nucleotide Archive in 2022. Nucleic Acids Res. 51, D121–D125 (2023).

59. Richardson, L. et al. MGnify: the microbiome sequence data analysis resource in 2023. Nucleic Acids Res. 51, D753–D759 (2023).

60. Schaaf, S., Erxleben-Eggenhofer, A. & Gruening, B. Galaxy and RDM: Being More Than a Workflow Manager: Living the Data Life Cycle. Proc. Conf. Res. Data Infrastruct. 1, (2023).

61. Batut, B. et al. Community-Driven Data Analysis Training for Biology. Cell Syst. 6, 752–758.e1 (2018).

62. Hiltemann, S. et al. Galaxy Training: A powerful framework for teaching! *PLOS Comput*. Biol. 19, e1010752 (2023).

63. Rasche, H. et al. Training Infrastructure as a Service. GigaScience 12, giad048 (2023).

64. Zierep, P., et al. How to increase the findability, visibility, and impact of Galaxy tools for your scientific community. Preprint at 10.37044/osf.io/qjbxc (2024).

65. Batut, B., et al. How to improve the annotation of Galaxy resources? Outcomes of an online hackathon for improving the annotation of Galaxy resources for microbial data resources. Preprint at 10.37044/osf.io/s7tru (2024).

66. Batut, B. et al. Galaxy CoDex for finding tools, workflows, and training. in F1000Research vol. 13 (2024).

67. Ison, J., et al. EDAM: an ontology of bioinformatics operations, types of data and identifiers, topics and formats. Bioinformatics 29, 1325–1332 (2013).

68. Royaux, C. et al. Guidance framework to apply best practices in ecological data analysis: lessons learned from building Galaxy-Ecology. GigaScience 14, giae122 (2025).

69. Eren, A. M. et al. Community-led, integrated, reproducible multi-omics with anvi’o. Nat. Microbiol. 6, 3–6 (2020).

70. Hadfield, J. et al. Nextstrain: real-time tracking of pathogen evolution. Bioinformatics 34, 4121–4123 (2018).

71. Maier, W. et al. Ready-to-use public infrastructure for global SARS-CoV-2 monitoring. Nat. Biotechnol. 39, 1178–1179 (2021).

72. Bazant, W., Blevins, A. S., Crouch, K. & Beiting, D. P. Improved eukaryotic detection compatible with large-scale automated analysis of metagenomes. Microbiome 11, 72 (2023).

73. Price, G. R. et al. Community-Curated Galaxy Interfaces with the Galaxy Labs Engine. Preprint at 10.20944/preprints202508.0199.v1 (2025).

74. Blankenberg, D. et al. Dissemination of scientific software with Galaxy ToolShed. Genome Biol. 15, 403 (2014).

75. Bray, S. et al. The Planemo toolkit for developing, deploying, and executing scientific data analyses in Galaxy and beyond. Genome Res. 33, 261–268 (2023).

76. The Bioconda Team et al. Bioconda: sustainable and comprehensive software distribution for the life sciences. Nat. Methods 15, 475–476 (2018).

77. Da Veiga Leprevost, F., et al. BioContainers: an open-source and community-driven framework for software standardization. Bioinformatics 33, 2580–2582 (2017).

78. Ison, J. et al. Tools and data services registry: a community effort to document bioinformatics resources. Nucleic Acids Res. 44, D38–D47 (2016).

79. Ison, J. et al. The bio.tools registry of software tools and data resources for the life sciences. Genome Biol. 20, 164 (2019).

80. Ponto, J. Understanding and Evaluating Survey Research. J. Adv. Pract. Oncol. 6, 168–171 (2015).

81. Cholewiak, S., Ipeirotis, P., Silva, V. & Kannawadi, A. scholarly. Zenodo 10.5281/ZENODO.7542349 (2023).

82. Kinney, R., et al. The Semantic Scholar Open Data Platform. Preprint at 10.48550/ARXIV.2301.10140 (2023).

83. Nasr, E., Pechlivanis, N., Zierep, P. & Batut, B. usegalaxy-eu/microgalaxy_paper_2025: v1.0.0. Zenodo 10.5281/zenodo.15088383 (2025).

