## Supplementary Document 1 for "Microbiology Galaxy Lab: The first community-driven gateway for reproducible and FAIR analysis of microbial data"

### Evaluation of the current usage and future needs of Galaxy for microbial data analysis

**Galaxy** is an **open-access, web-based platform for data-intensive computational research**. It is used in a variety of domains including **microbial data analysis**: for microbiome samples or bacterial isolates, long reads or short, shotgun or 16S, genomics, transcriptomics, or proteomics.

**microGalaxy**, a community of practice focused on microbial data analysis in Galaxy, would like to **learn** more about **use cases** where Galaxy was beneficial, and also the barriers **that prevent researchers from using Galaxy**. The aim is to collectively **find solutions** to improve user experience and reduce barriers**.** To this end, we are launching a **survey** to reach out to, and learn from, the microbial research community.

We encourage anyone to fill out this survey who:

- has used Galaxy for any kind of microbial focused research (e.g. metagenomics, metatranscriptomics, microbiome, bacterial isolates … ), or
- conducts microbial research or is planning to do so, but has not yet used Galaxy, or is hindered by certain aspects of Galaxy from using it.

You can find more information about this survey and how we plan to use the information collected on the dedicated post here:
https://galaxyproject.org/news/2023-03-21-microbial-survey/

Thank you for your help supporting better microbial data analysis in Galaxy!

#### Can you give a short description of  is your research area?

E.g.: I investigate the microorganisms in marine habitats to try and understand how they adapt to changing climatic conditions.)

<Free text>

#### What is your main target?

<One answer to select>

- Virus
- Bacteria
- Archaea
- Eukaryotes
- Mixed Communities

#### Which techniques do you use?

<Several answers can be selected>

- Single organism genomics
- Single organism transcriptomics
- Amplicon / metabarcoding
- Metagenomics
- Transcriptomics or metatranscriptomics
- Metabolomics
- Proteomics or metaproteomics
- Others

#### Which analysis do you use or would like to do?

<Several answers can be selected>

- Taxonomic profiling
- Pathogen tracking
- Assembly
- Antimicrobial resistance (AMR) detection
- Functional analysis
- Gene identification
- Single nucleotide polymorphism (SNP) identification
- Metagenome assembled genome (MAG) building
- Predictive metagenomics
- Comparative analyses
- Combination of multiomic approaches
- Phylogenetic tree construction
- Other

#### Which tools, platforms or databases are you using to conduct your research analyses

(e.g. Anvi'o, RStudio, nr, etc.)?

<Free text>

#### If the tools, platforms or databases from the previous question could be deployed in Galaxy would use prefer to use in there ?

<One answer to select>

- Yes
- No

#### How often do you use Galaxy?

<One answer to select>

- Never
- Less than once per year
- Several times per year
- Monthly
- Weekly
- Daily

#### If you never or rarely use Galaxy, what are the reasons?

<Free text>

#### Which Galaxy server(s) do you use?

<Several answers can be selected>

- Main, usegalaxy.org
- Galaxy Europe, usegalaxy.eu
- Galaxy Australia, usegalaxy.org.au
- Galaxy France, usegalaxy.fr
- Galaxy Norway, usegalaxy.no
- Galaxy Africa, africa.usegalaxy.eu
- Galaxy India, india.usegalaxy.eu
- Local Galaxy instance
- Other

#### Which tools and databases do you think are missing in Galaxy?

<Free text>

#### What features, tools, workflows etc. would you like to have in Galaxy that would support your research?

<Free text>

#### Which training resources are you using?

<Free text>

#### Which country are you based in?

<Free text>

#### What is your career stage?

<One answer to select>

- Undergraduate
- Graduate student

- PhD student
- Postdoctoral researcher
- Principal Investigator
- Staff scientist
- Other

#### Would you be interested in being contacted by the microGalaxy team and contributing an example use case to a manuscript?

<One answer to select>

- Yes
- No

#### If yes, please provide an email address

<Free text>

#### Do you have link(s) to publications or websites where we can read more about your research? If so, please add these below.

<Free text>
