## Supplementary Document 2 for "Microbiology Galaxy Lab: The first community-driven gateway for reproducible and FAIR analysis of microbial data"

### Microbial data analysis in Galaxy

Case studies

*You filled out our form "Evaluation of uses and needs for microbial data analysis in Galaxy". Thanks to you and the community, we collected **more than 100 answers**, which has given us valuable information about the community.*

*In your survey answer, you showed interest in being contacted by the microGalaxy team and contributing an example use case to the planned manuscript. You already gave us some basic information in the form, but **we would appreciate a more detailed description of your work**. We therefore request that you fill out the fields below **before August 31st**, so that we have a good overview of your use case.*

*We plan to **review and group case studies by topics,** and then **highlight especially intriguing ones** (after discussion with the community and you) in the main text of the paper. **All submitted studies (including yours) will be included in the Supplementary material,** and we invite the authors of the selected studies to become **coauthors of the manuscript.***

*Thank you for your help!*

#### Contact

| First name | Last name | Email | Affiliation | Country | ORCID |
|---|---|---|---|---|---|

#### Case study

**Title**

**Status**
*Possible answers: Starting / On-going / Completed / Other*

**Publications**

**Funding**

**Keyword**
*Give 3-8 keywords describing your use case*

### Research question

**Main target**
*Keywords. E.g. AMR, Pathogen, Microbiome community, MAGs building, Functional analysis, Gene identification, Comparative analysis, SNP analysis, or anything else*

**Short description of the research questions of your case study**
*Images can also be included*

#### Methods

**Experimental or Measurement technique used**
*E.g. metagenomics, metabarcoding, amplicon, proteomics, metabolomics, transcriptomics, single organism genomics*

**Data generation approach**
*E.g. Illumina, Nanopore, LS/MS*

**Data analysis approach**

*E.g. quality control, taxonomy profiling, assembly, differential gene expression, statistical analysis*

**Tools, workflows or platforms outside Galaxy**

**Galaxy tools used**

**Workflow(s) used**

**Specific benefits of using Galaxy**

*E.g. access to computational power, access to databases, tool availability, easy-to-use workflows, reproducibility, bulk processing using collections, or anything else*

**Short description of the methods you used**

*Images can also be included*

#### Interesting results

**Short description of the results you got**
*Images can also be included*

#### Other

**Anything else you would like to mention about your use case?**

### References

### Teaser of survey's results

*If you are interested to have a first peek at the results of the survey, here are some figures based on the first 90 answers.*

*The figure below shows the main techniques used by the participants.*

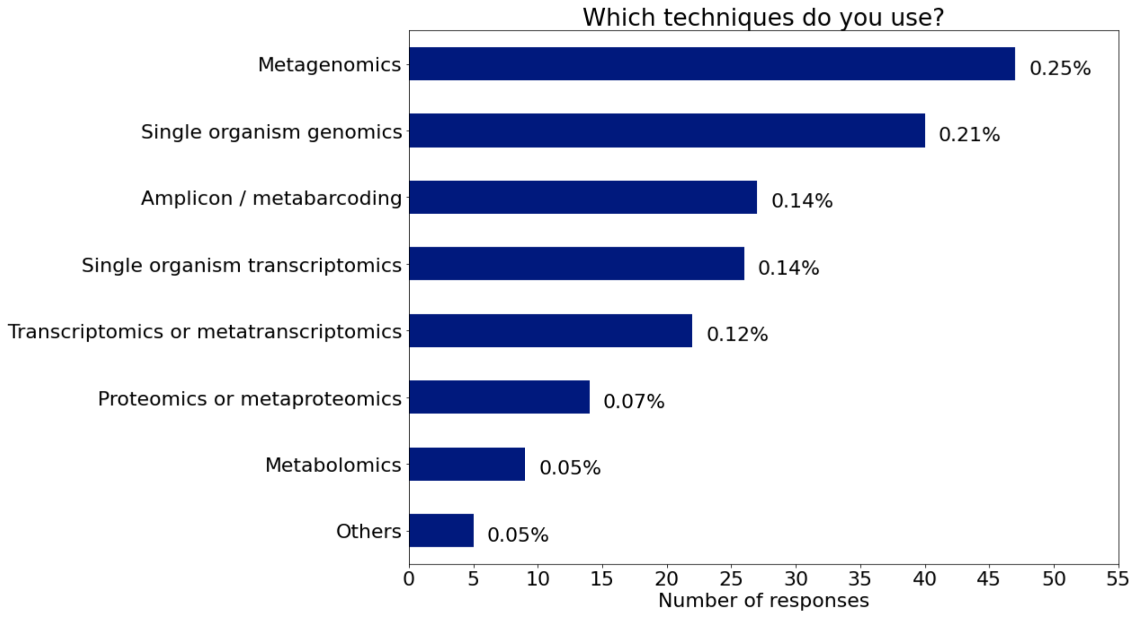

*We also discovered that a large majority of the respondents would use Galaxy for their analyses if the features they require were available on the platform. This motivates us to write this paper and take the next steps towards improving Galaxy for the microbial analysis community.*

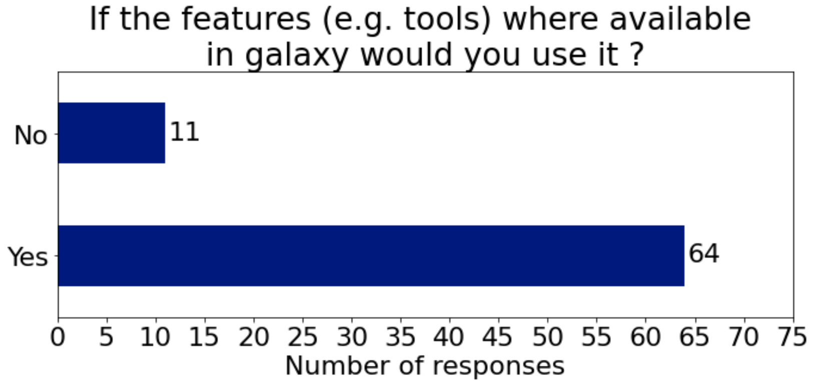
